# Sustained liver HBsAg loss and clonal T and B cell expansion upon therapeutic DNA vaccination require low HBsAg levels

**DOI:** 10.1101/2023.09.04.556204

**Authors:** Nádia Conceição-Neto, Wim Pierson, Maurizio Vacca, Matthias Beyens, Ben De Clerck, Liese Aerts, Birgit Voeten, Dorien De Pooter, Lore Verschueren, Koen Dockx, Mathias Vandenberk, Ewoud De Troyer, Kato Verwilt, Carl Van Hove, Mieke Verslegers, Leslie Bosseler, Marjolein Crabbe, Vinod Krishna, Isabel Nájera, Ellen Van Gulck

**Author notes:** **Corresponding author** Address correspondence to Ellen Van Gulck. **Funding sources:** Research was sponsored by Janssen Research and Development (Johnson & Johnson). **Conflict of interest:** Authors that are employees of Janssen Research and Development, may be Johnson & Johnson stockholders. **Financial support statement:** Research was sponsored by Janssen Research and Development (Johnson & Johnson). **Data availability statement:** All data associated with this study are present within the main text or the Supplementary Materials. **Authors’ contributions:** E.VG designed the studies. B.D.C, M.V, L.A, B.V performed the in vivo work. L.V and M.V performed and analysed the viral and blood peripheral read outs data. W.P, D.D.P, K.D setup, performed and analysed immunology related data. M.V performed Core and S staining’s of liver slides. L.B analysed the liver slides for HBV core and s-expression. K.V and C.V.H performed the single-cell RNA-sequencing experiments. N.CN, M.B and V.K analysed the single-cell RNA-sequencing and single-cell V(D)J data. N.CN, E.DT and M.C ran statistical analysis. N.CN, I.N and E.VG wrote the manuscript. All authors revised and approved the final manuscript.

## Abstract

**Background & Aims:** Suppression of HBV DNA, inhibition of HBsAg production and therapeutic vaccination to reverse HBV-specific T-cell exhaustion in chronic HBV patients are likely required to achieve functional cure. In the AAV-HBV mouse model, therapeutic vaccination can be effective in clearing HBsAg when hepatitis B surface (HBsAg) levels are low. The factor(s) required for mounting an effective immune control of HBV infection are unclear. Using a single-cell approach, we investigated the liver immune environment in the context of different levels of HBsAg as well as upon sustained HBsAg loss through treatment with an HBV specific GalNAc-siRNA followed by therapeutic vaccination.

**Methods:** C57BL/6 mice were transduced with a range of rAAV-HBV DNA to express different HBsAg levels. Mice were treated with GalNAc-siRNA targeting HBV transcripts to lower the HBsAg levels and then vaccinated 4 times with a DNA vaccine encoding HBV Core, Pol and Surface. We used single-cell RNA-sequencing on homogenised liver resident cells, paired with single-cell V(D)J receptor sequencing to understand the changes in the liver immune microenvironment.

**Results:** Treatment with GalNAc-HBV siRNA followed by therapeutic vaccination, achieved a sustained HBsAg loss in all mice. This was accompanied by an induction of CD4 follicular helper T-cell responses, polyclonal activation of CD8 T-cells in the liver and clonal expansion of plasma cells that were responsible for antibody production.

**Conclusions:** This study provides novel insight into the immune changes in the liver at the single-cell level, highlighting the correlation between the induced reduction in HBsAg levels and the clonal expansion of CD4 follicular helper T-cells, CD8 cytotoxic T-cells, plasma cells, and ISG-producing neutrophils in the liver upon HBV siRNA and subsequent therapeutic vaccine treatment.

**Lay Summary:** Chronic hepatitis B infection is characterized by a complex interplay between immune responses and viral replication in the liver. To achieve functional cure a combination of different treatments is likely required. In this study single-cell approach was used to understand the liver microenvironment in the context of different HBsAg levels followed by therapeutic vaccination in AAV-HBV mouse model and to identify key factors required to achieve functional cure.

**Graphical abstract:** 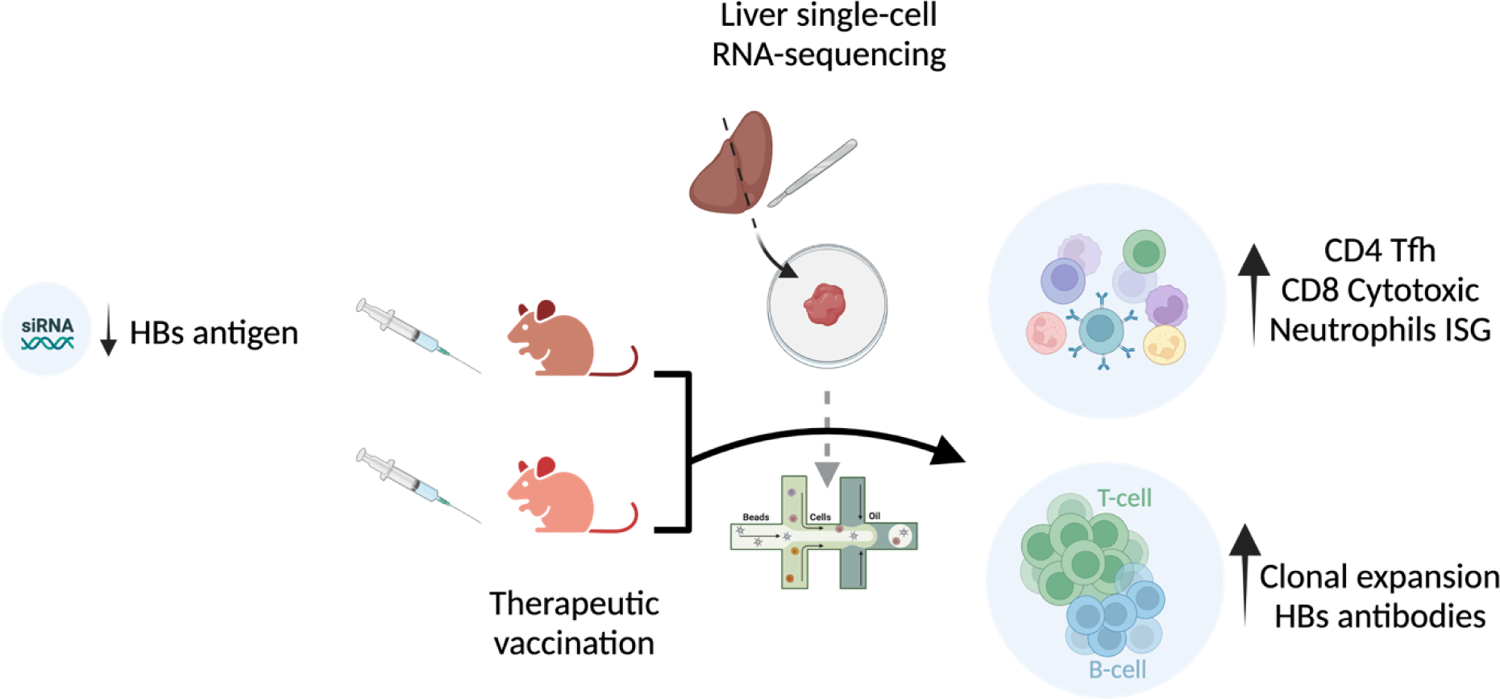

**Highlights:**

- AAV-HBV transduced mice sequentially treated with GalNAc-siRNA and therapeutic vaccine showed sustained HBsAg loss.
- The sustained HBsAg loss correlates with increased proportion and clonal expansion of CD4 follicular helper T-cells, CD8 cytotoxic T-cells, plasma cells, and ISG producing neutrophils in the liver.
- Baseline levels of HBsAg are important to determine outcome of therapeutic vaccination in mice.

## Introduction

The hepatitis B virus (HBV) poses a major global health problem, with the World Health Organization (WHO) estimating 1.5 million new infections yearly and nearly 296 million people living with chronic hepatitis B (CHB) infection, which is often life-long (1). Approximately 20-30% of individuals with CHB develop cirrhosis, liver failure or hepatocellular carcinoma (1). CHB has various clinical stages defined by HBV DNA levels, presence or absence of hepatitis B e antigen (HBeAg) and the presence or absence of liver inflammation defined by liver transaminase levels (ALT) (1).

Current standard-of-care is antiviral nucleos(t)ide analogue (NUC) therapy, that suppresses viral replication by inhibiting reverse transcription, but does not target the nuclear covalently closed circular DNA (cccDNA), therefore resulting in the persistence of HBV (2). Given that viral replication is supressed, and it largely eliminates the risk of liver cirrhosis but only partially the risk of hepatocellular carcinoma, it tends to be a lifelong therapy, since upon withdrawal, viral load usually rebounds, driven by cccDNA (3). For the subset of patients who can stop NUC therapy (HBeAg-, undetectable HBV DNA and low HBsAg) <3% achieve functional cure, through an unknown mechanism of action but with the key contribution of the HBV-specific adaptive immune response as observed in those who recover from acute infection. Used less often than NUC is finite treatment (typically 48 weeks) with pegylated IFN-alpha (+/-NUC), which can induce loss of HBV surface antigen (HBsAg) in up to 10 % of patients (4, 5). However, its low tolerability due to the well-known side-effects of IFN-based treatments can lead to early treatment discontinuation (6).

The current goal in HBV therapy is to achieve functional cure, defined as sustained loss of HBsAg (with or without HBsAg seroconversion) and undetectable HBV DNA in serum 24 weeks off treatment (7), where a functional HBV-specific adaptive immune response can control the infection (6). How HBV establishes a chronic infection in those that progress to CHB remains unclear but in acute HBV infection HBV-specific CD4, CD8 T-cells and neutralizing antibodies are key to limit the infection. Spontaneous immune control has been described in around 0.5% of chronically infected patients and is also accompanied by neutralizing antibodies and HBV-specific CD4 and CD8 T-cells (8). However, HBV persistence is characterized by lack of neutralizing antibodies, dysfunctional, exhausted HBV-specific T- and B-cells, inhibition and exhaustion of natural killer (NK) cells (9, 10). Although there is no definitive evidence of the role of HBsAg in T-cell dysfunction or exhaustion, a current hypothesis is that reduction of HBsAg levels is not sufficient but necessary to facilitate or mediate the gain in T-cell function to reach functional cure (11).

High levels of HBsAg (due to overproduction by a factor of 4 to 5log_10_ of subviral particles over infectious Dane particles) (12) are believed to be one of the mechanisms contributing to the sustained suppression of HBV specific immune responses, which hampers therapeutic vaccination (13). This hypothesis is also supported by the observation in animal models of CHB, which show that therapeutic vaccination is more effective when HBsAg levels are low (below 100 IU/ml) and it is not effective when HBsAg levels are >1000 IU/mL (14). In clinical setting studies, CHB infected individuals that received VPT-300; a prime-boost regimen, whereby the immune system is primed with an adenovirus (ChAdOx1) and boosted with a pox virus (MVA), showed significant and durable reductions of HBsAg only in those patients with baseline HBsAg levels <50IU/ml (15).

To better understand the liver immune microenvironment in HBV chronically infected patients, fine needle aspirates from patients at different stages of disease have been profiled using single-cell RNA-sequencing (16–19). Nkongolo *et al* profiled in depth the T-cell response in patients undergoing antiviral therapy and observed that CD8 hepatotoxic T-cells led to nonspecific hepatocyte killing, leading to fibrosis and cirrhosis (17).

In this paper we aimed to further understand the role of HBV antigenemia (i.e., HBs- and HBeAg levels) on the liver immune environment, to define key CD4 and CD8 T-cells populations and their ability to control the infection upon vaccination. Using single-cell RNA, T-cell receptor (TCR) and B-cell receptor (BCR) variability, diversity, and joining (V(D)J) sequencing of liver resident cells from (un)treated AAV-HBV infected mice were performed. We provide important insights into the changes in the liver microenvironment upon and during vaccination and in the context of different HBV levels.

## Methods

### Ethical Statement and Animal Experimentation

Animal studies were conducted in strict accordance with guidelines established by the Janssen Pharmaceutica N.V. Institutional Animal Care and Use Committee and following the guidelines of the European Community Council directive of November 24, 1986 (Declaration of Helsinki 86/609/EEC). The local Johnson and Johnson Ethical Committee approved all experimental protocols performed at Janssen. The criteria of the American Chemical Society ethical guidelines for the publication of research were met. Every effort was made to minimize animal discomfort and to limit the number of animals used. Mice were kept in a specific pathogen-free facility under appropriate biosafety level following institutional guidelines.

### Compounds

Therapeutic vaccine consisting of pDF-core, pDF-pol (20) and pDK-s plasmids. GalNAc conjugated HBV siRNA (GalNAc-HBV siRNA) and GalNAc-conjugated control siRNA (GalNAc-control siRNA) were manufactured by Arrowhead Pharmaceuticals and Axolabs respectively for research purpose only.

### Animal models and study designs

Female C57BL/6 mice (6-8 weeks old, Janvier labs, Le Genest-Saint-Isle, France) were transduced with different viral genome equivalents of rAAV-HBV1.3-mer WT replicon (BrainVTA, Wuhan, China) via the tail vein: for the low HBsAg level, 3×10^8^ viral genome equivalent (vge)/mice, for the mid-level 1×10^9^ vge/mice and for the high HBsAg level 2.5×10^9^ vge/mice. Treatment was initiated once persistent HBV viremia was reached (day 28 after transduction). Blood sampling was performed 8 days before treatment to randomize treatment groups based on HBsAg levels.

Mice were immunized intramuscularly (IM) in the anterior tibialis by electroporation (TriGrid electroporation system, Ichor Medical Systems, San Diego, CA, USA) with HBV vaccine containing 10µg of each plasmid (pDF-core, pDF-pol, pDK-s, 30µg in total, n=14) or with 30µg of empty plasmid (referred to as mock, only the group transduced with 3×10^8^ vge/mice). A vaccine boost was provided 3 weeks after prime with the same vaccine. Vaccinated mice (n=14) were split into 2 groups: one group received 2 vaccine doses; the other group received 4 vaccine doses. One week after the second vaccination (n=8) or 5 weeks after the last vaccination (n=6) splenocytes and intrahepatic lymphocytes were harvested and assessed for HBV-specific T-cell responses using IFN-γ ELISPOT (Fig. 1A). Blood for viral parameters was collected via the lateral saphenous vein every week, from which serum was prepared and stored at −80°C until assayed. At the end of the study the liver was formalin fixed and embedded in paraffin to determine HBs and HB core antigen levels.

**Fig. 1.**
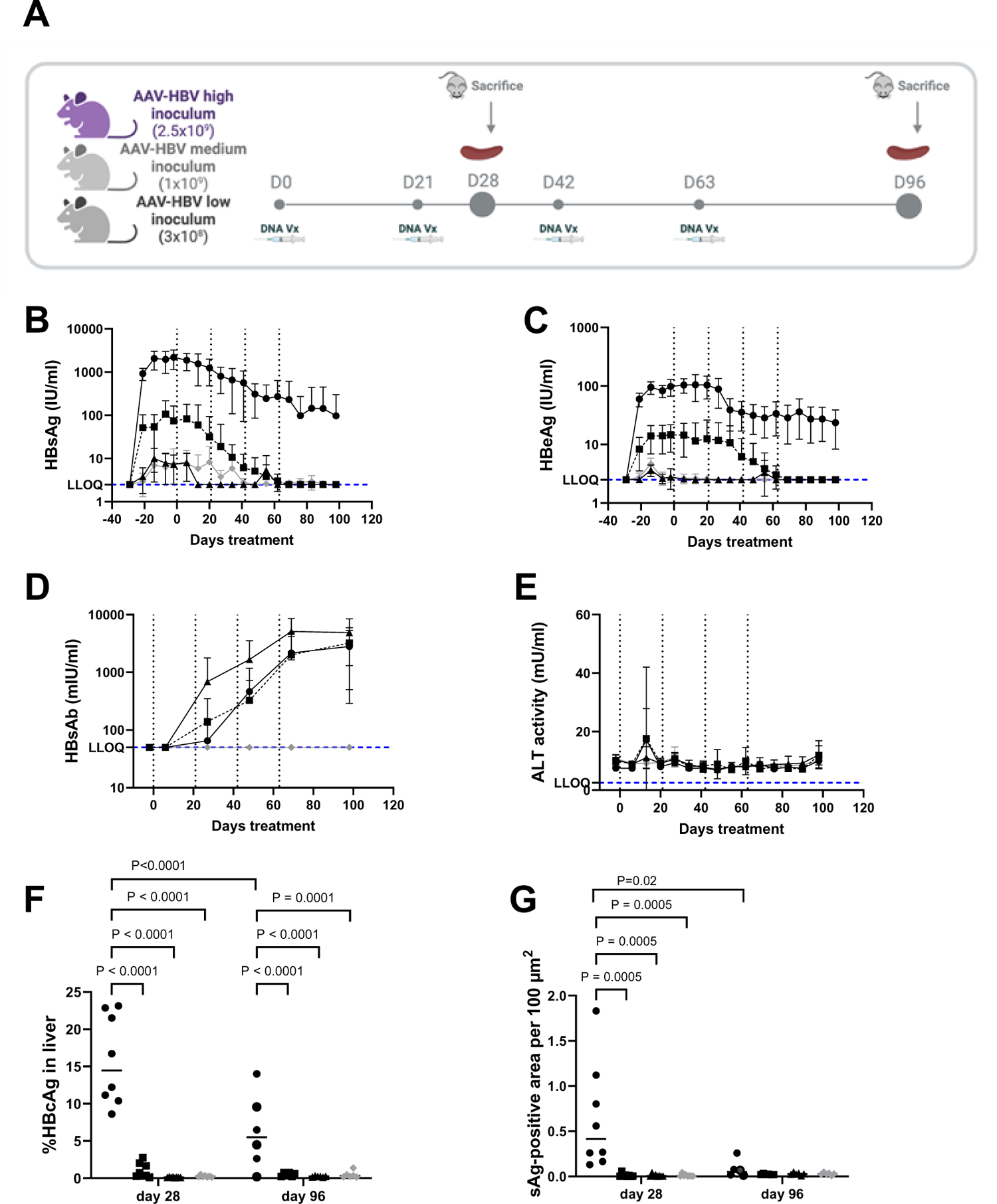
Effect of HBV levels on the efficacy of therapeutic vaccination in AAV-HBV mouse model. Female C57/Bl6 mice were transduced with different titers of AAV-HBV **(A)** (filled black circles symbols 2.5×10^9^vge, filled black square 1×10^9^vge and filled black triangle 3×10^8^vge) (Figure made with Biorender). 28 days after infection, treatment start with therapeutic vaccine encoding Core, Pol and Surface. Mice transduced with 3×10^8^vge were also treated with mock plasmid as a control (grey diamonds). Mice were vaccinated every 3 weeks for 4 times (indicated with dotted vertical lines). Every week serum samples were collected. Serum HBsAg **(B)**, HBeAg **(C)**, HBsAb **(D)** and ALT **(E)** were measured over time. Eight mice were sampled 1 week after second vaccine (28 days after treatment) and 6 mice 5 weeks after last vaccine (96 days after treatment). Data are presented as median +/- SD. FFPE livers were used to perform immunohistochemical stains for HB-core **(F)** and s **(G)** levels detection. Data were analyzed using mixed-effect analysis and Tukey’s multiple comparison testing (p values are indicated, otherwise it is not significant (ns)). Lower limit of quantification (LLOQ) of each assay is indicated with blue horizontal dotted line.

In a second study, female C57BL/6 mice were transduced with 2.5×10^9^ vge per mice of rAAV-HBV1.3-mer WT replicon to reach HBsAg levels of 3log_10_ IU/ml. At day 28 after transduction, mice were treated subcutaneously with 3mg/kg GalNAc-HBV siRNA (n=16) or with control GalNAc-siRNA (n=16) for 3 times on 3-weekly basis. At the third siRNA dose, mice were also treated with 30µg of the therapeutic vaccine or mock vaccine which was subsequently dosed 4 times every 3 weeks (Fig. 3A). HBV-specific T-cells were measured 1 week after the second vaccination (n=8/group) and 7 weeks after the last vaccination (n=8/group). Blood for viral parameters was collected every week, from which serum was prepared and stored at −80°C until assayed. At the end of the study, the liver was formalin fixed and embedded in paraffin to determine HBs and HB core antigen levels by immunohistochemistry. Liver resident T-cells were also collected to perform single-cell RNA-sequencing and V(D)J.

### Isolation of splenocytes and intrahepatic lymphocytes (IHL)

Spleens were isolated and tissue dissociation was performed using a GentleMACS Octo Dissociator (Miltenyi Biotec, Bergisch Gladbach, Germany) or by pressing through a 100 µm nylon cell strainer (Falcon, Corning, NY), and cells were added to coated ELISpot plates (Mabtech, Nacka Strand, Sweden). IHLs were obtained by perfusing the liver with PBS via the hepatic portal vein to flush out excess of blood. Ex vivo perfusion, enzymatic digestion and tissue dissociation of the left lateral liver lobe was performed using a GentleMACS Octo Dissociator with heaters and the liver perfusion kit for mice according to the manufacturer’s protocol (Miltenyi Biotec). Hepatocytes were separated from lymphocytes by centrifugation at 50× *g* for 5 minutes. Supernatant was spun down at 400× *g* for 5 minutes followed by resuspension in a 33.75% (v/v) Percoll (GE Healthcare) diluted in PBS with 2% foetal calf serum and density gradient centrifugation at 700 x *g* for 12 minutes. Next, residual hepatocytes and debris were discarded, and red blood cells co-sedimented with the intrahepatic immune cells (IHICs) were lysed using ACK lysis buffer (Lonza) for 5 minutes. Cells were washed twice and counted. Cell concentration and viability were determined using a Nexcelom Cellaca MX Cell Counter. Splenocytes were tested at 200,000 cells/well and IHLs were tested at 80,000 cells/well on IFN-γ ELISpot plates.

### Detection of HBV-specific T-cells by ELISPOT

All cells were stimulated with overlapping peptide pools covering the entire Core, Pol and S sequences (JPT, Gladbach, Germany) on pre-coated ELISpot plates (Mabtech, Nacka Strand, Sweden) to measure the number of cells that secrete IFN-γ. As a negative control, cells were stimulated with dimethyl sulfoxide (DMSO). After overnight stimulation, plates were developed following the manufacturers’ instructions (Mabtech, Sweden). The number of spot forming cells (SFC) was counted using an ELISpot reader (CTL, Cleveland, OH, USA). The DMSO (mock control) background was subtracted from all responses. Immunogenicity results are presented as the number of SFC per million of splenocytes or IHL. Results are shown as mean ± standard deviation. Due to the large size of the Pol protein, peptides were split into two peptide pools, Pol1 and Pol2.

### Viral parameters and alanine aminotransferase (ALT) analyses

Serum HBsAg, HBeAg and anti-HBs levels were quantified using CLIA kits (Ig Biotechnology, Burlingame, CA, USA). All CLIA kits were used according to the manufacturer’s guidance. Depending on the estimated levels, different dilutions of serum in PBS were used. Read-out of plates was performed with a Viewlux ultra HTS microplate imager (Perkin Elmer, Mechelen, Belgium).

The serum alanine aminotransferase (ALT) activity was analysed using a commercially available kit according to the manufacturer’s guidance (Sigma-Aldrich) and a Spark multimode microplate reader (Tecan, Mechelen, Belgium). Four µl of serum were used to perform the assay.

### Histology and Immunohistochemistry (IHC)

Formalin-fixed paraffin-embedded livers were sectioned at 5µm thickness, mounted on pre-charged slides and stained with HBcAg (B0586, Dako, Glostrup, Denmark) and HBsAg (ab859, Abcam, Cambridge, UK). Tissues were subjected to an EDTA-based antigen retrieval while signal amplification and detection was achieved with a hapten multimer and a DAB chromogenic detection kit, on the Ventana Discovery Ultra autostainer (Roche Diagnostics, Rotkreuz-Switzerland). Negative control staining was performed for every antibody using an isotype control (Rabbit PE IgG for HBcAg, mouse IgG1 for HBsAg). Sections were coverslipped with the automated Ventana HE600 (Roche Diagnostics) and scanned using the Hamamatsu Nanozoomer RX.

### Image analysis

IHC stainings were evaluated using the HALO^TM^ system (Halo 3.4). Liver tissue was discriminated from glass/lumen by using a tailored random forest classifier in HALO v3.4. and expression of HBcAg and HBsAg were evaluated using two tailored algorithms based on nuclear or signal area detection, respectively. Output parameters include percentage of HBcAg-positive cells, relative to total number of haematoxylin-positive cells and HBsAg-positive surface area, relative to the total classified liver area analysed.

### Single-cell RNA-sequencing and data analysis

IHL’s were processed according to 10x Chromium Single Cell V(D)J Reagent Kits User Guide and loaded on the Chromium 10x platform using the 5’ v1.1 chemistry (10x Genomics). Libraries were sequenced on a NovaSeq4000 (Illumina) to an average ∼50,000 reads per cell. Read alignment was done using the Cell Ranger pipeline against the mouse genome reference (mm10). Resultant cell by gene matrices for each sample were merged across all conditions tested and samples. Pre-processing, alignment, and data filtering was applied equivalently to all samples using OpenPipeline v0.7.0 workflows (github.com/openpipelines-bio). Cells with less than 1000 UMIs or less than 200 genes or more than 25% mitochondrial counts were removed from downstream analysis. All downstream analysis was done in R using the Seurat v4.1.1 package (21).

Data was log-normalized with a scaling factor of 10,000. For the first level clustering, the top 2,000 most variable genes were selected (‘vst’ method implemented in FindVariableFeatures()) and scaled using a linear model in the ScaleData() function. Afterwards, principal component analysis (PCA) was run, the number of significant principal components (PCs) to be used for downstream cell clustering was determined using an ElbowPlot and heatmap inspection. A nearest neighbour graph and Uniform Manifold Approximation and Projection (UMAP) plot were generated using the significant PCs.

A Louvain clustering was run on all cells, and the best resolution for clustering was determined using an average silhouette scoring across all clusters, testing 10 resolutions between 0.1 and 1 as previously implemented in Ziegler et. *al* (22). Marker genes for each cluster were calculated using the FindAllMarkers() function (method= ‘wilcox’) implemented in Seurat and each cluster was iteratively subclustered further using the same approach. Sub-clustering was stopped when the resulting clusters were not meaningfully different. Clusters were annotated as cell type populations based on the expression of genes that are known markers of specific cells by expert annotation and using the mouse liver atlas (23).

### Single-cell V(D)J analysis

V(D)J (TCR and BCR) sequencing data was processed using the Cell Ranger V(D)J pipeline with mm10 as a reference. All downstream analysis was done in Python using the *scirpy* v0.12.2 library (24) and the *dandelion* v0.3.2 library (25), respectively for TCR and BCR analysis. TCR and BCR annotations were further refined via V/D/J reannotation of constant region using IMGT’s mouse reference v3.1.38 (26), and only sequences corresponding to CH1 region for each constant gene/allele were retained. For the contigs assembled, low-confidence, non-productive or with UMIs<2 were discarded. If five or more cells had identical dominant α–β pairs, along with no more than 15% mismatches in the CDR3 nucleotide sequences, the dominant α–β pairs were identified as clonal TCRs. Only cells that were annotated as T-cells with the cell typing were considered for this analysis. Similarly for B cells, a threshold of 3 cells was defined for clonality of BCRs. TCR and BCR sequencing data was processed using the Cell Ranger vdj pipeline with mm10 as a refence. For more in depth BCR sequencing analysis, data obtained from the Cell Ranger pipeline was analysed using the *Immcantation* toolkit (27, 28). Specifically, igblast V(D)J gene assignments were generated for each sample using the AssignGenes.py script, and a database for each sample was generated using MakeDb.py. Consequent to this, each sample was filtered to remove unproductive sequences and cells with multiple heavy chains using the *Changeo* R package and cells were split by their heavy and light chains. For the samples, pairwise Hamming distances were estimated. We found that some samples had few clonal expansions resulting in a small threshold distance cut-off. Consequently, we chose a manual threshold of 0.12 as the distance cut-off and combined the samples together for further clonal analysis. Hierarchical clustering was performed for the sequences that were within this distance threshold and lineages were built using “buildPhylipLineage”. Finally, clonal trees and somatic hypermutation frequencies were inferred using the “observedMutations” command from the *shazam* R package, and trees were constructed using plotTrees in the *alakazam* R package. An R markdown script that contains the details of this analysis is available in the supplementary information.

### Statistical analysis

Statistical comparisons were performed using either GraphPad Prism 9 or the package rstatix v0.7.2 in R. Statistic differences between the different treatment groups on immune responses and expression of HB antigen in liver was calculated using mixed-effect analysis and Tukey’s multiple comparison testing. Single-cell data differential proportion was calculated using a pairwise Wilcoxon test and FDR multiplicity adjustment (Benjamini-Hochberg) was done for all comparisons per each cell population. All p-values <0.05 were considered significant. Due to small sample size, both raw and adjusted p-values are reported.

## Results

### Low baseline HBsAg levels are determinant for the therapeutic vaccination efficacy

The AAV-HBV mouse model of chronic HBV infection, shown to recapitulate the HBV-specific immune tolerance observed in CHB patients during long (months) HBV replication F(19, 29), was used to evaluate the effect of HBV DNA therapeutic vaccination.

To evaluate the potential immunosuppressive effect of HBV antigens, namely HBsAg and HBeAg, mice were transduced with increasing amounts of AAV-HBV DNA (more specifically 3×10^8^, 1×10^9^ and 2.5×10^9^ vge/mice, Fig. 1A) to achieve different stable levels of HBsAg. Four weeks after transduction, and once HBV antigen levels where stable, mice were vaccinated with a mixture of three plasmids (pDNA) each encoding for Core, Pol or S antigens by intramuscular administration with electroporation at week 0, 3, 6 and 9. The control mice were transduced with AAV-HBV DNA (3×10^8^vge/mice) and electroporated with mock vaccine at the same days. The therapeutic vaccine was able to clear antigenemia (reduce HBV antigens (HBsAg and HBeAg) levels to undetectable) after a single (prime) vaccination in the low (<10IU/ml) HBsAg group at baseline, and after 4 vaccinations for the mid (100IU/ml) HBsAg group (Fig. 1B-C, Fig. S1-S2). All control mice transduced with lowest vge/mice cleared antigenemia naturally, but this happened only at a later time point (by day 60 after start treatment). For those with higher baseline levels of HBsAg (Fig. 1B-C, Fig. S3), sustained loss of HBsAg levels were achieved in 50% of mice despite the induction of HBsAb (Fig. 1D) and only 1 mouse showed sustained HBeAg loss (Fig. S3).

ALT levels were transiently increased in some mice after the first vaccination (Fig. 1E). HBsAb were induced in all groups after a second vaccination. Mice with lower baseline HBsAg levels (10IU/ml) induced more rapidly and higher HBsAb levels compared to those that started treatment with HBsAg baseline levels of 100 or 1000 IU/ml (Fig. 1D).

At day 28 (1 week after second vaccination) and at day 96 (5 weeks after last vaccination), liver HBcore (HBcAg)- and HBs-antigens were measured by immunohistochemistry (Fig. 1F-G). The lower the baseline HBsAg levels, the lower the levels of liver HBcAg and HBsAg were at day 28. The number of cells in the liver that expressed Core and S significantly decreased over time from day 28 to day 96, respectively (p<0.0001 for HBcAg, p=0.02 for HBsAg) for the group starting with high baseline HBsAg levels (1000IU/ml). For the group with about 100 IU/ml at baseline, the decline observed in Core levels was not significant over time, due to the low levels already reached upon 2 vaccinations.

### HBsAg baseline levels in the peripheral circulation inversely correlate with HBV-specific immune response upon vaccination

To further investigate the mechanism of HBV antigen clearance upon vaccination, and the role of HBV antigenemia in immune tolerance, immune responses were evaluated 1 week after second vaccination (Fig. 2A) and at 5 weeks after last vaccination across the low, middle and high baseline HBV antigen levels (Fig. 2B). HBV-specific IFN-γ responses were induced against all tested HBV-antigens (Core, Pol and S). Interestingly, the induced immune response against Core and S measured 1 week after second vaccination (Fig. 2C), as well as 5 weeks after last vaccination (Fig. 2D), inversely correlated with the serum HBsAg level at the start of the treatment. The induced responses to Pol remained comparable irrespective of the serum HBsAg baseline level (Fig. 2E-F). This inverse correlation was also observed in terms of HBsAb responses, with higher level of HBsAb observed in the low baseline HBsAg level group (Fig. 2G). Five weeks after last vaccination, besides splenocytes, also intrahepatic lymphocytes (IHL) were evaluated. Responses in the liver were higher compared to spleen for Core (p<0.0001). T-cell responses to Pol and HBsAg were comparable between liver and spleen (Fig. 2H). This demonstrates that in AAV-HBV infected mice, 4 weeks after transduction, HBsAg levels have an impact on the induced immune responses.

**Fig. 2.**
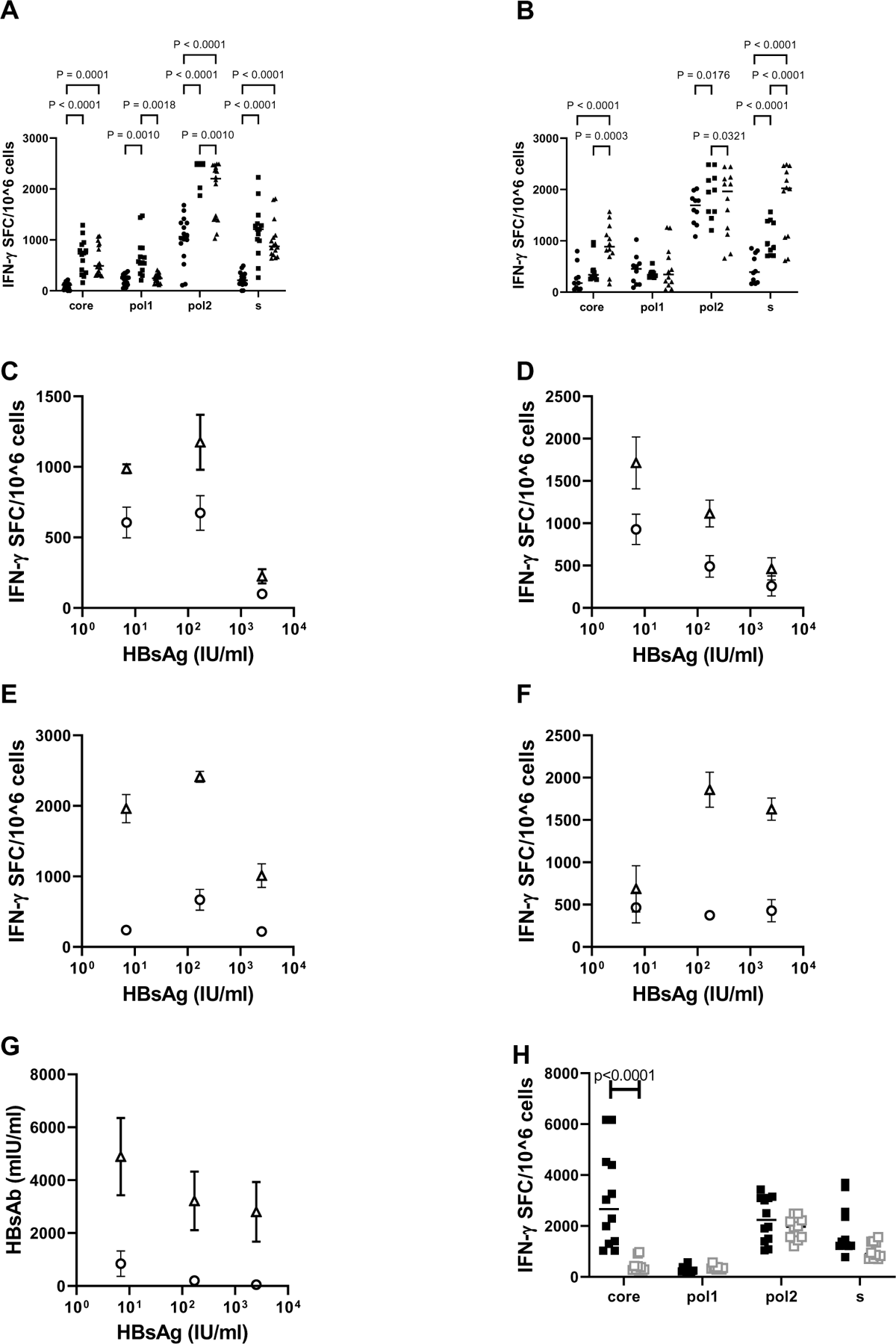
Induction of HBV-specific immune responses in AAV-HBV mice treated with therapeutic vaccine. As mentioned in Fig. 1 part of the mouse liver samples were collected at day 28, is 1 week after second vaccine (**A**) and the other part at the end of the study day 96, is 5 weeks after last vaccine (**B**). Splenocytes were stimulated directly with Core, Pol or Surface peptide pools in IFN-γ ELISpot plates. IFN-γ spot forming cells (SFC) per million of splenocytes are plotted on the y-axis. (Filled black circles symbols 2.5×10^9^vge, filled black square 1×10^9^vge and filled black triangle 3×10^8^vge). Data are DMSO subtracted and mean of triplicate is shown for each mouse. p-values represent differences between the different groups in inducing immune responses against certain peptide pool, they are calculated using mixed effect analysis and Tukey’s multiple comparison testing. In panels **(C and D)** the mean SFC and standard deviation are plotted against the mean HBsAg levels measured in serum at start of the treatment (triangle are s-specific immune responses, circles are Core specific responses). Showing data for 1 week after second vaccine and 5 weeks after last vaccine respectively. In panels (**E and F**) the mean SFC and standard deviation are plotted against the mean HBsAg levels measured in serum at start of the treatment (circles are Pol1 and triangles are Pol2 specific responses). Showing data for 1 week after second vaccine and 5 weeks after last vaccine respectively. **(G)** Correlation between induction of HBs antibodies and HBsAg levels at start of treatment (open circles HBsAb responses at 1 week after second vaccine, open triangles responses at 5 weeks after last vaccination). **(H)** At the end of the study intrahepatic lymphocytes are isolated from each mouse and directly stimulated with the different peptide pools in ELISpot assay. In this figure intrahepatic responses (filled black squares) are compared with responses induced by splenocytes (open grey square) from the same mice.

### Can HBV tolerance be broken by HBV siRNA treatment prior to therapeutic vaccination?

To mirror the levels of HBsAg in CHB patients, we used the AAV-HBV mouse model with baseline HBsAg levels of 1000IU/ml. Using this setting, we investigated whether treatment with siRNA to lower HBsAg and HBeAg levels for 6 weeks prior to vaccination could improve vaccine efficacy levels. Four weeks after AAV-HBV transduction, once HBsAg levels were stable, mice were treated with 3 doses of GalNAc-HBV siRNA (n=32) or control GalNAc-siRNA (n=32) at 3-week intervals, followed by 2, or 4 doses (n=16 active and 16 control siRNA per regimen) of pDNA vaccine at 3-week intervals starting on week 10 post transduction (Fig. 3A).

**Fig. 3.**
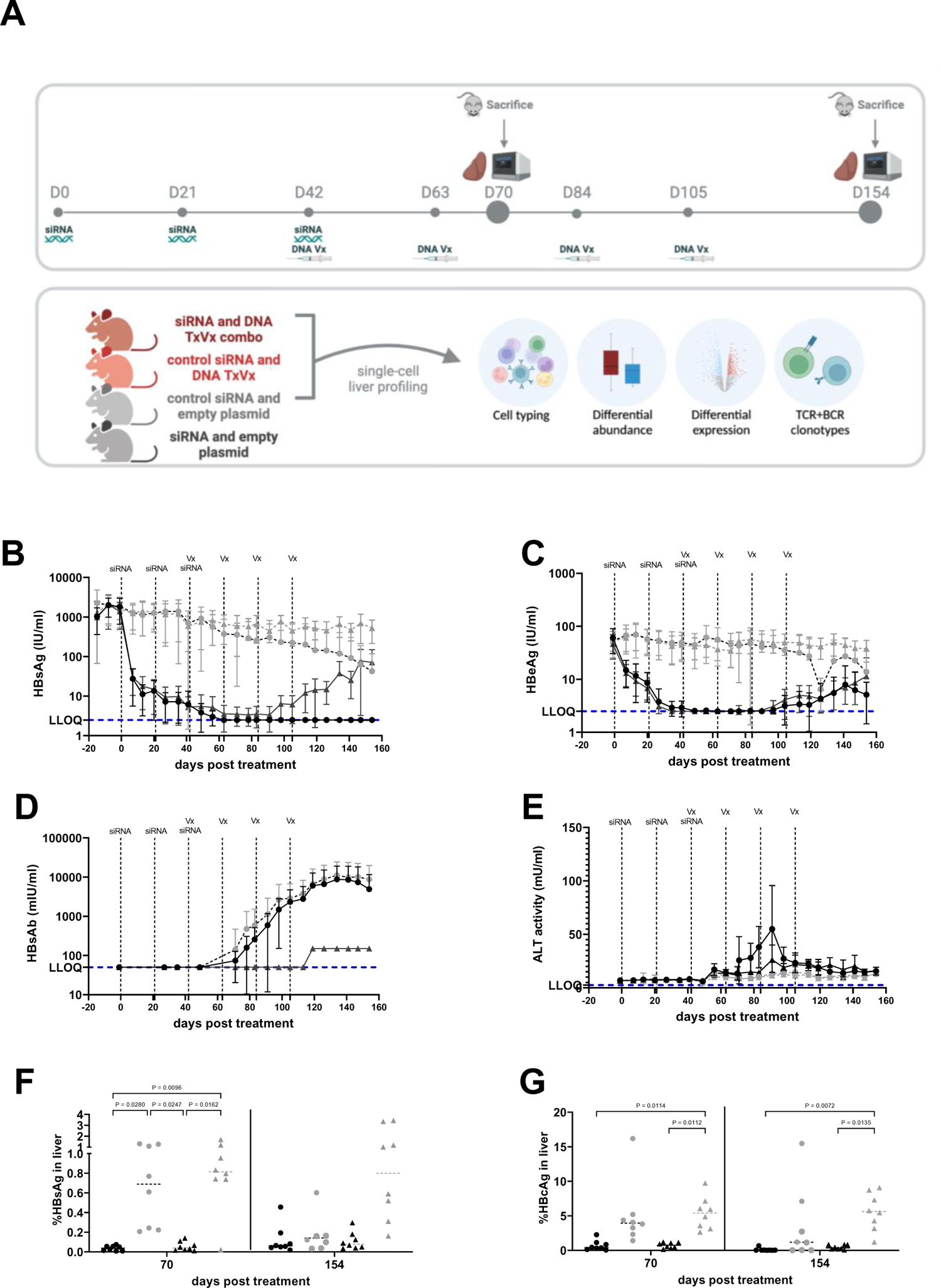
Effect on HBV viral parameters by sequential treatment of siRNA and therapeutic vaccination in AAV-HBV mice. Female C57/Bl6 were transduced with 2.5×10^9^vge of AAV-HBV **(A)** 4 weeks after infection, treatment started with siRNA and followed by therapeutic vaccine encoding Core, Pol and Surface-antigens. Mice were vaccinated every 3 weeks (indicated with dotted vertical lines and Vx) for 4 times after 3 doses of siRNA (indicated with dotted vertical lines and siRNA) (Figure made with Biorender). Every week, blood was taken from each mouse and serum was collected for measurement of HBsAg **(B)**, HBeAg **(C)**, HBsAb **(D)** and ALT **(E)** over time. Eight mice were sampled at day 70 (1week after second vaccination), the other mice were samples at day 154 (7 weeks after last vaccination). Immunohistochemistry on FFPE livers was used to semi quantitatively assess HBsAg **(F)** and HB core Ag **(G)**. Mixed effect analysis and Tukey’s multiple comparison testing were performed as statistical testing (p values are indicated, otherwise it is not significant). Black circles: GalNAc-HBV siRNA and TxVx; black triangles: GalNAc-HBV siRNA and mock vaccine, grey circle: GalNAc-control siRNA and TxVx, grey triangles: GalNAc-control siRNA and mock vaccine. Lower limit of quantification (LLOQ) of each assay is indicated with horizontal blue dotted line.

Mice that received the control GalNAc-siRNA and the mock vaccine did not show any decline in viral parameters (Fig. 3B-C, Fig. S4). In contrast, GalNAc-HBV siRNA rapidly reduced serum HBsAg level by 2log_10_ IU/ml and HBeAg by >1log_10_ IU/ml (Fig. 3B-C, Fig. S5-S6). Animals that received GalNAc-control siRNA and then the therapeutic vaccine showed a moderate maximum decline of 1log_10_ in HBsAg (Fig. 3B-C, Fig. S7), comparable with results observed in AAV-HBV mice infected with 2.5×10^9^ vge/mice (Fig. 1B).

GalNAc-HBV siRNA treatment alone failed to trigger HBV-specific immunity, measured by HBsAb titers (Fig. 3D) and IFN-γ ELISpot for Core, Pol or S (Fig. 4) and consequently viral relapse was observed 6 weeks after the last administration of siRNA. In contrast, all mice treated with GalNAc-HBV siRNA and therapeutic vaccine showed stable long term (until day 154) loss of serum HBsAg, with development of high titers of anti-HBs (Fig. 3D) and induction of HBV-specific T-cells (Fig. 4), though loss of HBeAg was only observed in some of the animals (Fig. 3C, Fig. S5D). Moreover, the reduction of HBV antigenemia through the sequential treatment with GalNAc-HBV siRNA prior to therapeutic vaccination was well tolerated with no effects on body weight and only a moderate and transient ALT elevation detected after the first vaccine (<100mIU/ml, Fig. 3E).

**Fig. 4:**
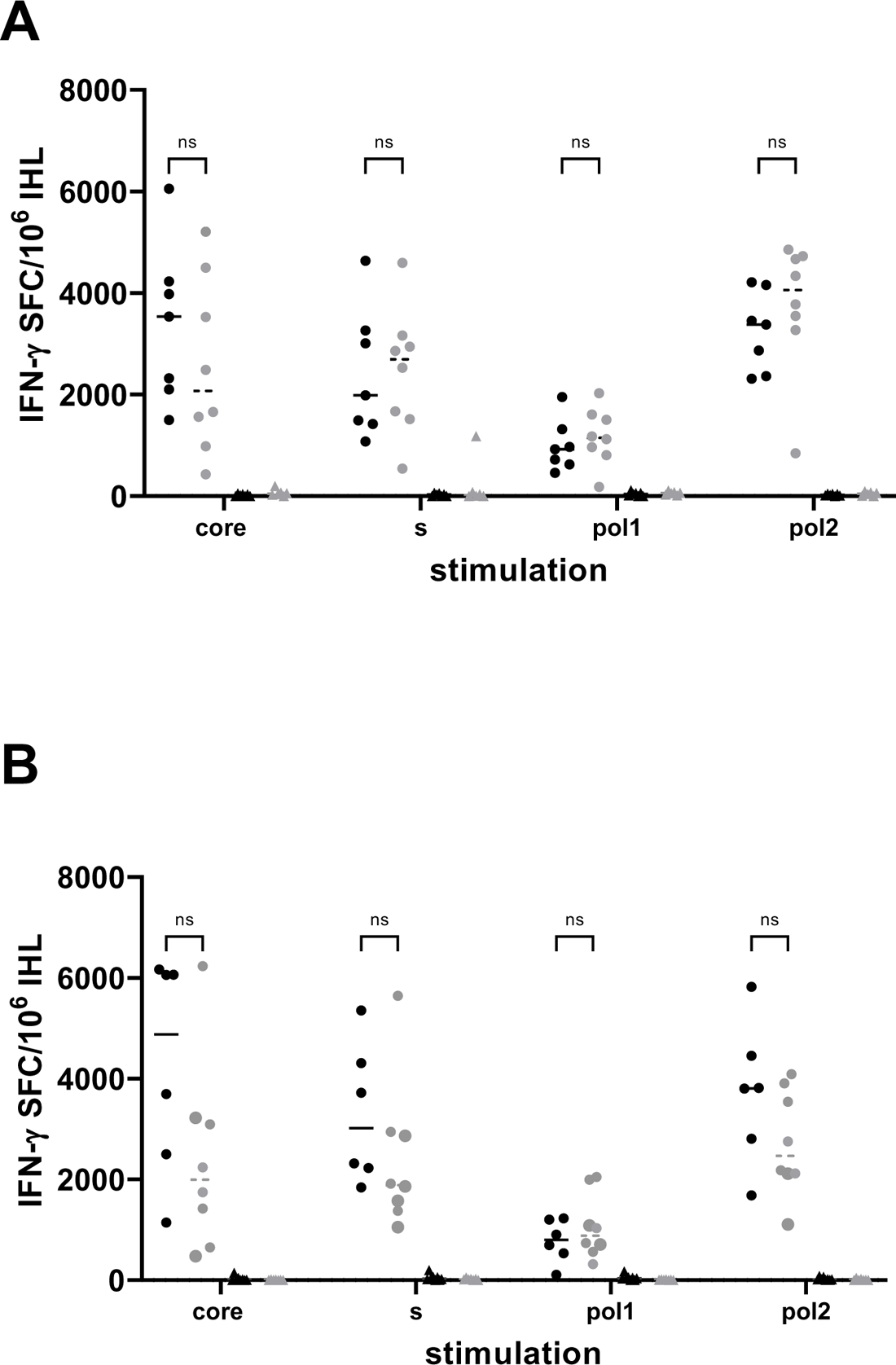
Induction of HBV-specific immune responses in AAV-HBV mice sequentially treated with GalNac-siRNA and therapeutic vaccine. IHL from day 70 (1week after second vaccination) **(A)** and from day 154 (7 weeks after last vaccination) **(B**) were stimulated directly ex vivo in IFN-γ ELISpot plates with Core, Pol, or Surface peptide pools. Results are presented on the y-axis as spot forming cells (SFC) per million of IHL, DMSO background is subtracted. Each dot or triangle is the mean of 3 replicates for each mouse. The horizontal line represents the median within one group. Black circles: GalNAc-HBV siRNA and TxVx, black triangles: GalNAc-HBV siRNA and mock vaccine, grey circle: GalNAc-control siRNA and TxVx, grey triangles: GalNAc-control siRNA and mock vaccine. P values are represented and calculated by Tukey’s multiple comparison test.

An important aspect of this study is the analyses of the liver immune environment in the context of different baseline levels of HBV and upon (or no) treatment. In concordance with the HBsAg serum levels data, the levels of HBsAg in the liver at 1 week after second vaccination (Day 70) were reduced upon treatment with GalNAc-HBV siRNA, irrespective of whether they were subsequently vaccinated or not (p=0.0096, p=0.0162 respectively, Fig. 3F). As described before, no reduction in HBsAg levels were observed in those mice groups treated with GalNAc-control siRNA, either vaccinated from week 6 or not.

At day 154 (7 weeks after last vaccination) the HBsAg levels in the liver remained low for the groups treated with GalNAc-HBV siRNA, irrespective of their vaccination status. Moreover, multiple vaccinations had also an effect on HBsAg in the liver, since mice treated with GalNAc-control siRNA and therapeutic vaccine showed a further decline in HBsAg. For HBcAg in the liver, a comparable profile was seen as described for HBsAg (Fig. 3G). At day 70, treatment with GalNAc-HBV siRNA had a significant decreasing effect on expression of HBcAg in the liver irrespective of vaccination. At day 154, the HBcAg levels remained low in the groups treated with GalNAc-HBV siRNA and those treated with control siRNA and therapeutic vaccine showed a decrease in HBcAg (Fig. 3G). These data corroborate the previous data that suggests that (higher) efficacy of therapeutic vaccination requires levels of HBsAg <100 IU/ml. We showed that an HBV siRNA is be an effective agent to reduce HBsAg levels and enables and improves the efficacy of the therapeutic vaccine.

To investigate and identify potential correlates of protection, the induction of liver HBV-specific T-cells was evaluated 1 week after second vaccination (Day 70, Fig. 4A) and 7 weeks after last vaccination (Day 154, Fig. 4B). HBV-specific T-cells, measured by IFN-γ ELISPOT, could be detected in the liver of all mice groups vaccinated with HBV-expressing pDNA. At day 154 after the start of treatment the strength of HBV-specific immune responses increased against all HBV antigens compared to responses at day 70. Taken together, these data show that treatment with GalNAc-siRNA prior to therapeutic vaccination enables high level of HBV-specific T-cell responses. Mice vaccinated with mock vaccine did not show any induction of HBV-specific T-cells irrespective of their treatment with GalNAc-siRNA.

### Liver CD4 Tfh-like cell subpopulation induced by vaccination identified by single-cell RNA-sequencing analysis

To identify which liver cells are most relevant to break T-cell tolerance, single-cell RNA-sequencing was performed to generate a liver cells atlas. Liver resident cells from three groups of mice transduced with 2.5×10^9^ vge of AAV-HBV and treated with either GalNAc-HBV siRNA and therapeutic vaccine (G1), GalNAc-control siRNA and therapeutic vaccine (G2) or control GalNAc-siRNA and mock vaccine (mock group, G4), were analysed using single-cell RNA-sequencing 1 week after second vaccine administration (day 70, first timepoint, Fig. 3A) and 7 weeks after the 4^th^ vaccine administration (day 154, second timepoint, Fig. 3A). A total of 4 samples per group per timepoint were profiled, except for the first timepoint of the GalNAc-HBV siRNA and therapeutic vaccine and the control groups (Fig. S8A). The uneven sample size between the first and second timepoint hampered a meaningful statistical comparison. After filtering for quality and doublet identification during cell typing, 250,966 cells across a total of 22 samples were obtained. Across all mouse liver samples, the lowest number of cells obtained was 8,590 and the maximum was 15,623 (Fig. S8A). After annotation, 11 major cell types were obtained: B cells, T-cells, cholangiocytes, dendritic cells, endothelial cells, granulocytes, hepatocytes, innate lymphoid cells (ILCs), monocytes, neutrophils and stromal cells. The majority of the cells detected across the full dataset were liver sinusoidal endothelial cells (LSECs) (27%) and hepatocytes (24%). B cells (8%), T-cells (16%) and monocytes (14%) were also well represented, and the remaining cell types made up a smaller fraction of the data (Fig. 5A; Table S1). The major subpopulations were further annotated to provide more granularity to each cell type (Fig. S8B), leading to a total of 39 subpopulations. Within the T-cell compartment, a total of 13 subpopulations (Fig. 5B) were identified, representing CD8, CD4 and NK-T cells. Of note, within the CD4 compartment, a subpopulation with phenotype of a T-follicular helper-like cell, expressing *Pdcd1* (PD-1), *Ifng*, *Cxcr3*, *Il21* and *Cd40lg* (Fig. 5C) was identified. This subpopulation was significantly increased (raw p-value=0.029, adjusted p-value= 0.044) in both vaccinated groups versus mock vaccinated group on day 154, though an increased trend at day 70 was already observed (Fig. 5D).

**Fig. 5:**
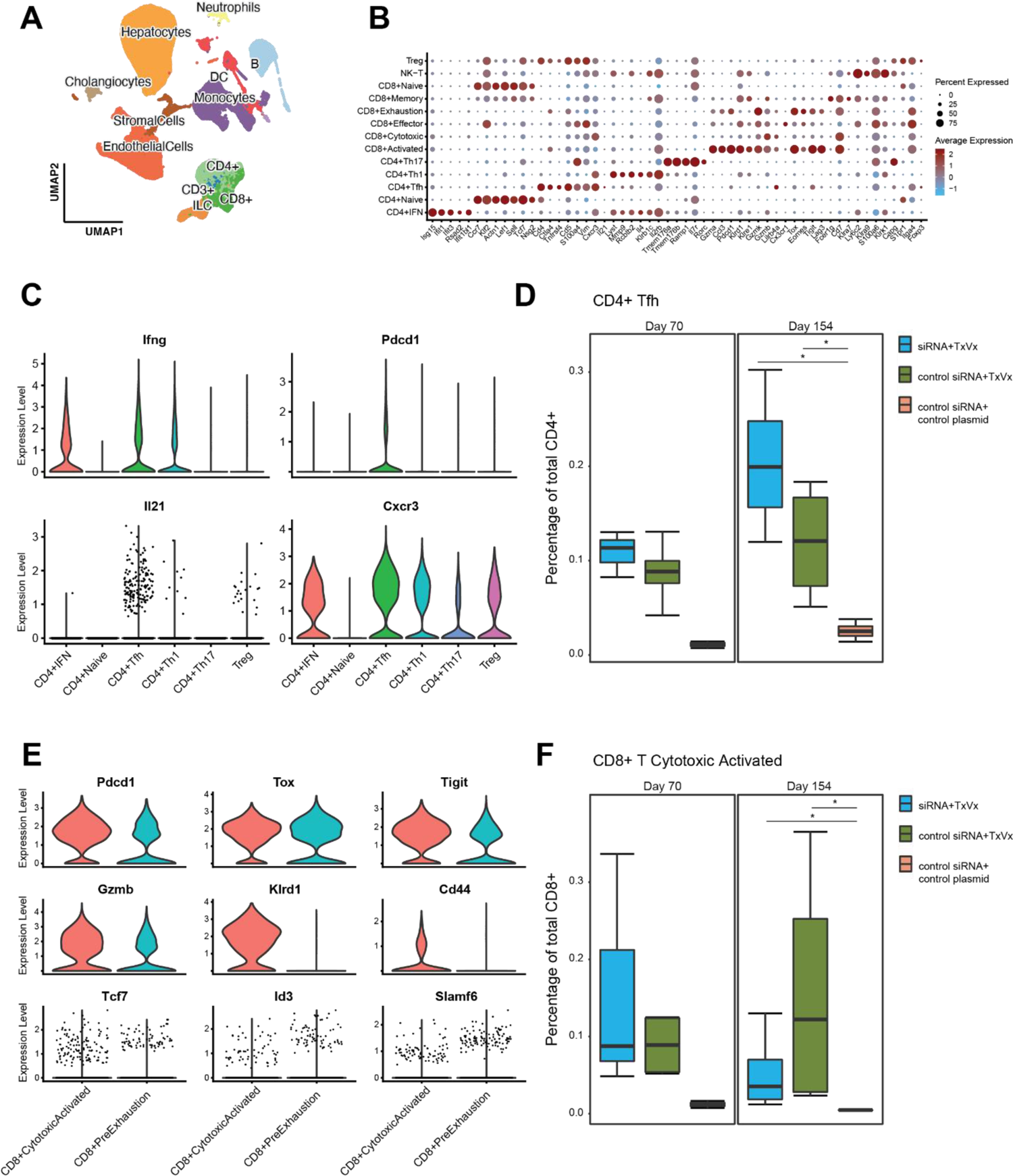
Single-cell liver atlas of the AAV-HBV model. **(A)** Overview of the full liver single-cell RNA-seq atlas using UMAP highlighting the top-level cell annotation. **(B)** Dot plot of top differentially expressed genes across the T-cell compartment. **(C)** Violin plot from CD4 Tfh genes across the CD4 T-cell compartment. **(D)** Box plot of CD4 Tfh cell percentages in total CD4 T-cells in the different groups faceted across the two sampling timepoints. **(E)** Violin plot from effector and exhaustion genes in CD8 cytotoxic activated and CD8 pre-exhaustion. **(F)** Box plot of CD8 cytotoxic activated cell percentages in total CD8 T-cells in the different groups faceted across the two sampling timepoints. Pairwise Wilcoxon test, with fdr correction. comparison testing (*p<0.05)

### Decrease in liver naïve CD8 T-cells and increase in pre-exhausted naïve cells upon vaccination

The CD8 compartment comprised of naïve, central memory, effector memory, cytotoxic T-cells and two populations expressing *Pdcd1* (PD-1), *Tox*, *Gzmk* and *Tigit*: cytotoxic activated and pre-exhausted T-cells (Fig 5B, 5E). The cytotoxic activated cells were distinguished from the pre-exhausted population based on their cytotoxic phenotype and their expression of *Cd44*, *Klrd1* and higher levels of *Gzmb* (Fig. 5E). This cytotoxic activated CD8 population and CD8 pre-exhausted T-cells were significantly increased upon vaccination irrespective of treatment with siRNA at day 154 versus the mock group (raw p-value=0.026, adjusted p-value= 0.039, Fig. 5F; Fig. S9A, raw p-value=0.029, adjusted p-value= n.s., respectively). In contrast, the central memory T-cell compartment showed a significantly decreased frequency in the vaccine treated groups, irrespective of treatment with siRNA, compared to mock treated group at day 154 (Fig. S9A; raw p-value=0.029, adjusted p-value= 0.044). Frequency comparison between the treatment groups showed a trend (albeit non-significant) of decreased naïve T-cells in both vaccinated groups irrespective of siRNA in comparison with the mock group (Fig. S9A). Finally, the cytotoxic T-cells showed increased frequency only in the group that received GalNAc-HBV siRNA and subsequent HBV DNA vaccination, but not in the GalNAc-control siRNA and therapeutic vaccine group compared to mock treated group at day 154 (Fig. S9A, raw p-value=0.029, adjusted p-value= n.s.).

To assess their proliferation capacity, we looked at *Tcf7* (TCF-1) and *Id3* expression (30, 31). Both populations (pre-exhausted and cytotoxic activated) expressed *Tcf7* in the treatment groups, albeit at low levels (Fig. 5E). In the pre-exhausted T-cells expression of *Tcf7* and *Id3* was higher in the vaccine groups compared to the mock vaccinated group (Fig. S9B).

### Macrophage, NK cell and neutrophil compartments change upon therapeutic vaccination

Within the monocyte and dendritic cell compartments, we identified 4 subtypes of dendritic cells (cDC1, cDC2, migratory DCs and pDCs as described in (23)), 3 subtypes of macrophages (Kupffer cells, capsule macrophages and macrophages) and 3 subtypes of monocytes (classical, patrolling and proliferating monocytes), which were annotated according to their hallmark genes in Fig. S10A). A higher frequency of macrophages was observed between the two vaccinated groups irrespective of siRNA treatment versus the mock group (Fig. S10B; raw p-value=0.029, adjusted p-value= 0.044) on the second timepoint sampled (day 154). No changes were observed in Kupffer cells.

As previously mentioned, even though the uneven sample size between the first and second timepoint hampered a statistical comparison, we observed that the frequency of cDC1s remained stable across the two timepoints, when GalNAc-HBV siRNA treatment was followed by therapeutic vaccination (G1). In contrast, both the mock group (G4) and the group that received control-siRNA and therapeutic vaccine (G2) showed a trend of increased frequency at day 154 compared to day 70 (Fig. S10C, raw p-value=0.029, adjusted p-value=0.087). The cDC2 population showed an increased trend at day 154 only in the GalNAc-HBV siRNA followed by therapeutic vaccine group (G1), whereas it remained comparable in the other groups. Across the three treatment arms, a decrease in frequency on day 154 was observed in the pDCs (Fig. S10C) compared to day 70. Migratory DCs at day 154 were significantly increased in GalNAc-HBV siRNA followed by vaccine group (G1) versus the mock vaccine group (G4) (Fig. S10C, raw p-value= 0.029, adjusted p-value =0.057).

In the NK compartment, the population CD11b+CD27-(high cytolytic function) was increased at day 154 in the groups not receiving GalNac-HBV siRNA (Fig. S11A; raw p-value=0.029, adjusted p-value= 0.087). This showed with an inverted trend with the CD11b-CD27+, with a cytokine producing phenotype. CD11b+CD27-cells expressed high levels of *Gzmb*, *Prf1*, *Nkg7*, whereas CD11b-CD27+ expressed *Cd160* and *Il2* (Fig. S11B).

The neutrophil compartment, despite having low numbers of cells, showed interesting trends. We were able to detect three populations: an immature population expressing *Ltf*, *Lcn2* and *Camp* and two mature populations (Fig. S12A). One of them expressing typical type I IFN signature genes such as *Isg15*, *Mme*, *Rsad2* and *Cd274* (PD-L1); the other expressing immunosuppressive markers like *Mmp8*, *Mmp9, Arg1* and *Padi4* (Fig. S12B). In addition, the mature populations also showed an interesting trend across the different arms, where the GalNAc-HBV siRNA followed by therapeutic vaccination group (G1) showed higher frequency of the type I IFN (ISG) neutrophils and lower immunosuppressive (MMP9) neutrophils than both the control siRNA and therapeutic vaccine groups (G2) and the mock group (G4) (Fig. S12C).

### Therapeutic vaccination induces TCR clonal expansion across multiple T-cell subtypes

To investigate whether our vaccination strategy induced clonality in the T-cell compartment, single-cell TCR sequencing was performed. The data-driven cell type annotation was paired with the TCR repertoire clonality for further granularity (Fig. S13A-B). The CD8 T-cell compartment showed clonal TCR expansion on both groups receiving vaccination irrespective of siRNA treatment. The pre-treatment with GalNAc-HBV siRNA and subsequent therapeutic vaccination group (G1) showed 33% of the CD8 T-cells with clonal expansion at the first timepoint (day 70), increasing to 53% on the second timepoint (day 154). The group treated with GalNAc-control siRNA followed by therapeutic vaccination (G2) showed 31% of the CD8 T-cells with clonal expansion at the first timepoint (day 70) and 39% on the second timepoint (day 154). For the mock group (G4) it was observed at 1 and 2%, respectively (Table S2).

When looking into each CD8 T-cell subtype, CD8 naïve and CD8 central memory T-cells did not show expansion across the different groups (Fig. S13B). Interestingly, the CD8 T-cell population with the cytotoxic activated phenotype (expressing *Pdcd1*, *Gzmb*, *Klrd1* and *Cd44*) showed over 70% of clonality in the groups that received the therapeutic vaccine irrespective of siRNA treatment and independent of the number of vaccines received (Fig. 6A). The pre-exhaustion population showed high clonality in the group that received GalNAc-HBV siRNA followed by therapeutic vaccine (G1), with 76% of cells clonally expanded in the first timepoint (day70) and 90% in the second timepoint (day 154). For the group treated by GalNAc-control siRNA followed by therapeutic vaccine (G2), the first timepoint showed 44% clonal expansion and increased to 73% on the second one (Fig. 6A).

**Fig. 6:**
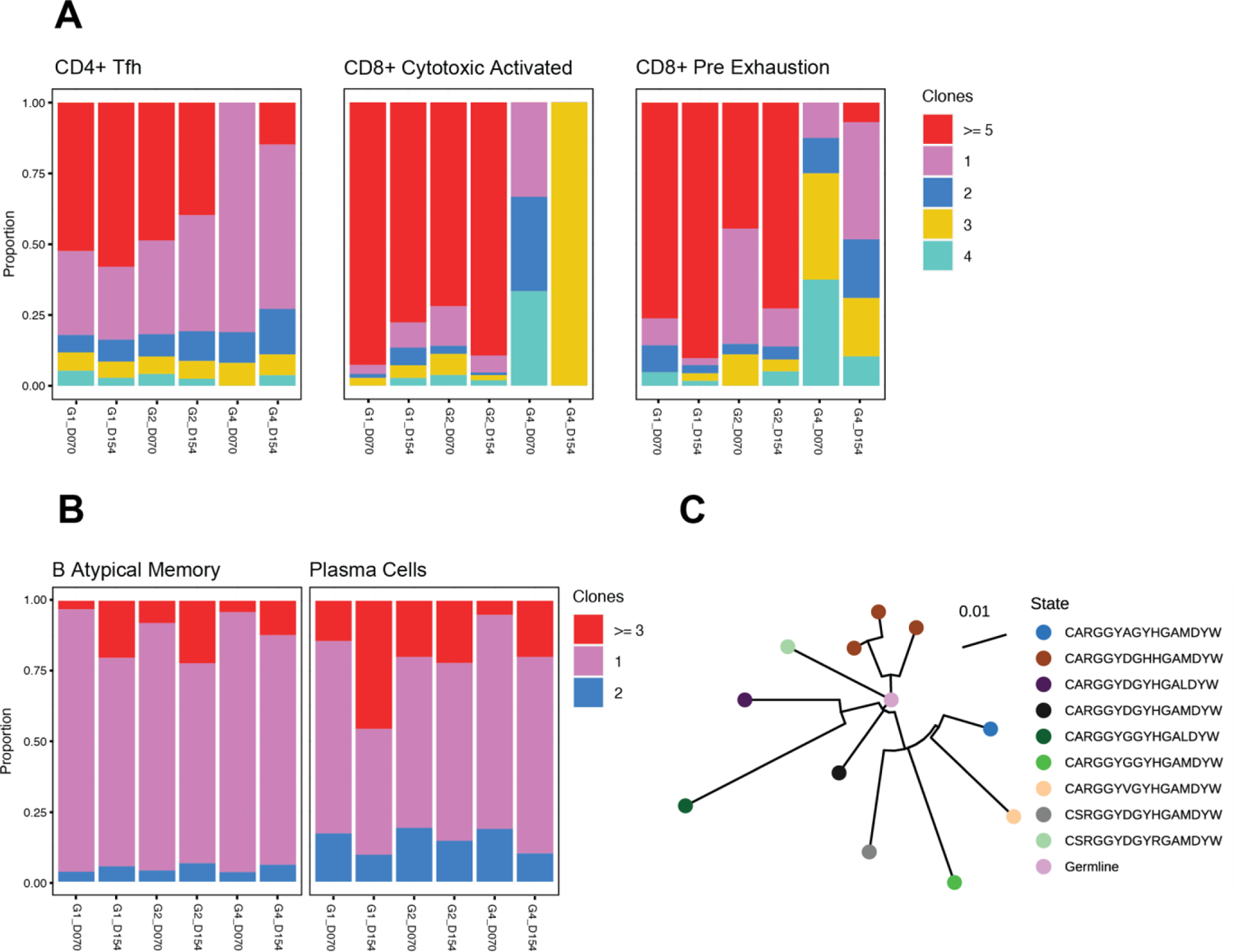
TCR and BCR clonality. **(A)** Proportion of CD4 Tfh, CD8 cytotoxic activated and CD8 pre-exhaustion with clonal expansion (represented by >=5 clones). **(B)** Proportion of B atypical and plasma cells with clonal expansion (represented by >=3 clones). **(C)** Clonal phylogenetic analysis from CDR3 sequences from a siRNA and therapeutic vaccine sample. G1 represents treatment with GalNAc-HBV siRNA followed by therapeutic vaccine, G2 represents GalNAc-control siRNA and therapeutic vaccine and G4 represents GalNAc-control siRNA and mock vaccine.

In the CD4 T-cell compartment, a more modest expansion was observed, and mainly driven by the CD4 Tfh cells (Fig. 6A, Fig. S13A). On a global CD4 T-cell level, the GalNAc-HBV siRNA followed by therapeutic vaccine group (G1) showed 8% clonal expansion at the first timepoint (day 70), and 23% second timepoint (day 154). The GalNAc-control siRNA group followed by therapeutic vaccine (G2) showed 9% on both timepoints. For the mock group (G3) it showed 3 and 2%, respectively (Table S2).

When breaking down the clonal expansion based on the transcriptional profile cell annotation, for the CD4 Tfh, the group treated with GalNAc-HBV siRNA and subsequent therapeutic vaccine (G1) remained stable with 52 and 58% clonal expansion at both timepoints. For the group treated with GalNAc-control siRNA followed by therapeutic vaccination (G2), clonality reached 49 and 40% on both timepoints, respectively. In the mock group (G4), no clonal expansion was detected on the first timepoint, and 15% on the second. Interestingly, the Treg compartment showed no clonal expansion in the mock group, while a 14% on the first timepoint and 12% clonality at the second timepoint were observed for the group that received GalNAc-HBV siRNA followed by therapeutic vaccination (G1). The group receiving GalNAc-control siRNA and therapeutic vaccine (G2) showed 5 and 7% clonality, respectively (Fig. S13A).

### BCR clonal expansion across atypical B cells and plasma cells

Four different B cell subpopulations were detected: activated B cells expressing *Myc*, *Nr4a1*, *Ier2* and *Nfkbid*, classical memory B-cells expressing *Ighd* and *Fcer2a*, atypical memory B cells expressing *Fcrl5* and *Lgals1* and finally plasma cells expressing *Jchain*, *Xbp1* and *Igha* (Fig. S14A). We observed BCR clonal expansion only in atypical memory B and plasma cells, but not in activated B-cells and classical memory B cells (Fig. 6B). Plasma cells sampled from mice receiving GalNAc-HBV siRNA followed by therapeutic vaccine (G1) showed a proportion increase in clonal expanded cells from 14% in the first timepoint (day 70) to 46% at day 154. The group receiving control siRNA followed by therapeutic vaccine (G2) showed 20% and 22% clonality in each timepoint, respectively. Within the atypical B cell subset, the second timepoint for both treatment groups reached 20% clonality (Fig. S14B).

To understand whether the clonally expanded plasma cells expressed long-lived plasma cell genes (32), we checked for *Sdc1* (encoding CD138), *Tnfrsf17* (encoding BCMA) and *Epcam* (encoding CD326). A lower expression of all genes was observed in plasma cells with only 1 clone (Fig. S14C). Correspondingly, no clonal expansions were detected in the absence of vaccination. All phylogenetic analyses are described in Fig. S14D-F.

## Discussion

Despite considerable scientific advancements in the field of HBV therapeutic vaccination, the key factors essential for breaking HBV tolerance continue to be elusive. The current study provides novel insights into the immunological effect that HBV levels have on the liver immune environment using the AAV-HBV mouse model in a setting mirroring the HBsAg level observed in CHB patients. Two approaches were taken: one was the use of the AAV-HBV model with low, middle and high levels of HBsAg. The second approach was to first lower HBsAg levels through treatment with an HBV-specific siRNA and follow with vaccination. Herein we describe single-cell analyses to study liver immune microenvironment dynamics when lowering HBsAg levels through treatment with GalNAc-HBV siRNA prior to therapeutic vaccination that provides insightful information to guide potential therapeutic approaches.

Using a pDNA-based vaccine encoding for Core, Pol and S antigens, we demonstrate that high-level HBsAg expression correlated with HBV-specific immune tolerance in AAV-HBV mice, preventing successful therapeutic vaccination and HBsAg clearance, as previously described (14). The inhibition of HBV antigen expression in hepatocytes using a liver-directed GalNAc-HBV siRNA enabled a therapeutic vaccine approach to elicit its full immunogenic capacity and led to sustained HBsAg loss in mice with HBsAg levels around 3log_10_ IU/ml or less (14). This was accompanied by the induction of both HBV-specific T- and B-cell responses, likely driving the sustained suppression of HBsAg. The induction of both HBV-specific CD4 and CD8 T-cells accompanied by neutralizing antibodies is also seen in the human setting with resolver patients (8).

The observed enhancement of HBsAb in AAV-HBV mice could be a concern, since the induction of anti-HBs could act as a decoy for measuring HBsAg in serum, which could bias its detection. To circumvent this, in this study HBsAg levels were measured in both serum and liver. Concomitant with the decline in the serum, HBsAg was also decreased in the liver of animals that received vaccine alone, clearly indicating a treatment effect. Furthermore, in the group that received the GalNAc-HBV siRNA followed by therapeutic vaccination, a further reduction on HB Core antigen in the liver was observed, which was accompanied by a transient and small ALT increase after the first vaccination. Taken together, these findings clearly indicate that there is a vaccine effect on HBV antigens likely by induction of T-cells driven by the HBsAg levels in serum and liver.

To further understand the immune microenvironment in the liver during vaccination, and whether prior GalNAc-HBV siRNA treatment impacted liver immune dynamics, single-cell RNA- and V(D)J-sequencing were performed. Though limited by small sample size, the high number of cells per sample allowed for an in-depth cell population analysis. Mice treated sequentially with GalNAc-HBV siRNA and therapeutic vaccine cleared HBV antigenemia, which correlated with both strong CD4 Tfh responses, TCR clonal expansions as well as with plasma cell clonal expansion. In contrast, mice treated with control siRNA (that therefore maintained high levels of HBV antigenemia) followed by therapeutic vaccination did not all cleared HBV antigenemia despite strong induction of CD4 Tfh and TCR clonal expansions. Given that the main difference with the group that cleared HBV was a more modest plasma cell clonal expansion, it suggests that mounting a CD4 Tfh response alone is not sufficient for clearing HBV and that induction of B-cells is a key factor (33).

It is well established that CD4 Tfh cells are key in supporting germinal centre B-cells to produce vaccine-specific immunoglobulins (34) and CD4 T-cell priming has been reported as critical for a successful therapeutic vaccination in AAV-HBV mice with a recombinant protein prime/MVA boost (35). Our study shows that anti-HBs were detectable in serum, starting at 70 days post treatment. CD4 Tfh cells were detected already on the first single-cell analysis timepoint (day 70) in both vaccinated groups, indicating that these were able to support B-cells mounting a productive antibody response. The B-cell compartment showed the highest clonal expansion at day 154, and especially within plasma cells of mice treated with both GalNAc-HBV siRNA and therapeutic vaccine. Correspondingly, upon phylogenetic reconstruction, no clonally expanded lineages were detected for B-cells for any of the treated groups at the early timepoint (day 70). For day 154 we found clonally expanded lineages, with greater clonal expansion in both groups that received the vaccine.

In the CD8 T-cell compartment, the cytotoxic activated population, expressing PD-1, as well as effector markers, was significantly increased in both vaccinated groups in comparison with the control group. Moreover, these showed pronounced clonal expansion, indicating a vaccination related activation. Although the differences between mice that received vaccine alone and those that received the sequential treatment of GalNAc-HBV siRNA and therapeutic vaccine were not significant, there was a clear trend of more TCR clonal expansion in the dual treatment group over time compared to the vaccine only group. Combining these data with the ELISpot data, leads us to suggest that induction of both T and B-cells is important, as well as the function and clonal expansion of these cells and represents a correlate of protection.

Besides effector markers, *Pdcd1 (PD-1) and Tox* were also upregulated on CD8 T-cells, which warrants further monitoring to understand whether these cells might progress to a terminal exhaustion phenotype, or whether this is a transient effect of T-cell activation. The cells expressing *Pdcd1* and *Tox* also expressed *Tcf7*, *Id3* and *Slamf6*, therefore were not terminally exhausted and indicate a potential transient state. A previous report from our group (19) has shown terminal exhaustion in AAV-HBV mice with high stable HBsAg levels (4log_10_IU/ml), therefore we postulate that the mice with lower stable HBsAg levels (2-3log_10_IU/ml) used in this study did not elicit a terminal exhausted phenotype (19). Nonetheless, the potential benefit of treatment with immune checkpoint inhibitors is being explored (36).

Lastly, in the neutrophil compartment, some interesting trends were observed. The mature neutrophil population MMP9, expressing immunosuppressive markers showed higher frequency in the control group and therapeutic vaccine alone group compared to the mice that received the sequential treatment. These neutrophils have been stated to have potent immunosuppressive capabilities in oncology (37). The same might be true in the context of chronic infection, where it has been described that HBV may reduce neutrophil responses (38). The more activated neutrophils (expressing *Isg15*) showed the opposite trend, with higher frequencies in the treated groups, which could indicate that these might be mediating infected hepatocyte killing. A similar neutrophil cell type expressing type I interferon activation has already been described in chronic hepatitis B patients (16), pointing towards possible translation in humans.

In conclusion, this study provides novel insight into the immune changes in the liver at the single-cell level, showing a correlation between the induced reduction in HBsAg levels and the clonal expansion of CD4 follicular helper T-cells, CD8 cytotoxic T-cells, plasma cells, and ISG-producing neutrophils in the liver upon HBV siRNA and subsequent therapeutic vaccine treatment. These factors and novel insights can help guide new immune approaches towards functional cure.

## Abbreviations

AAV: adeno-associated virus

BCR: B cell receptor

CDC: constitutive dendritic cell

CHB: chronic hepatitis

B GalNAc: N-acetylgalactosamine

HBV: hepatitis B virus

HbsAg: hepatitis B surface antigen

HbeAg: hepatitis B e antigen

HbsAb: hepatitis B surface antibody

HBcAg: hepatitis B Core

IHC: immunohistochemistry

IHL: intrahepatic lymphocytes

ISG: interferon stimulating genes

LSEC: Liver sinusoidal endothelial cells

MVA: modified vaccinia Ankara

NK-cell: natural killer cell

PDC: plasmacytoid dendritic cell

TCR: T-cell receptor

Tfh: T follicular helper cell

UMAP: Uniform Manifold Approximation and Projection

UMI: Unique Molecular Identifier

vge: viral genome equivalents

WHO: World Health Organization

## Acknowledgements

We would like to thank the following individuals for their expertise and assistance throughout all aspects of our study: Frederik Pauwels, Jan-Martin Berke, Richard May and Elien Peeters. We would like to thank Charles River laboratories (CRL) Beerse Belgium. We would like to thank Heather Davis for proofreading the manuscript for content and language.

