## supplemental figures for "Sustained liver HBsAg loss and clonal T and B cell expansion upon therapeutic DNA vaccination require low HBsAg levels"

**SUPLEMMENTARY Appendix:**

**Table of contents**

**Supplemental results:**

**Figure S1. Viral parameters over time of individual mice that were initially transduced with AAV-HBV ($3x10^8$ vge/mice)**

**Figure S2. Viral parameters over time of individual mice that were initially transduced with AAV-HBV ($1x10^9$vge/mice)**

**Figure S3. Viral parameters over time of individual mice that were initially transduced with AAV-HBV ($2.5x10^9$vge/mice)**

**Figure S4. Viral parameters over time of individual mice that were initially transduced with $2.5x10^9$vg/ml rAAV-HBV and treated with GalNAc-control siRNA and mock vaccine.**

**Figure S5. Viral parameters over time of individual mice that were initially transduced with AAV-HBV ($2.5x10^9$vg/ml) and treated with GalNAc-HBV siRNA and therapeutic vaccine (TxVx).**

**Figure S6. Viral parameters over time of individual mice that were initially transduced with $2.5x10^9$vg/ml rAAV-HBV and treated with GalNac-HBV siRNA and mock vaccine.**

**Figure S7. Viral parameters over time of individual mice that were initially transduced with $2.5x10^9$vg/ml rAAV-HBV and treated with GalNAc-control siRNA and therapeutic vaccine (TxVx).**

**Figure S8. Single-cell RNA sequencing QC and annotation.**

**Figure S9. CD8 T cell frequency and pre-exhaustion profiles.**

**Figure S10. Monocyte and DC compartment frequency analysis.**

**Figure S11. ILC compartment frequency analysis.**

**Figure S12. Neutrophil compartment analysis.**

**Figure S13. TCR clonality frequency.**

**Figure S14. B cell compartment analysis.**

**Supplementary Table 1 – Percentage distribution of cell types**

**Figure S1- Viral parameters over time of individual mice that were initially transduced with AAV-HBV (3x10[8] vge/mice).** Individual-level data are shown; data corresponds to the mean data plotted in Figure 1. (A) Hepatitis B surface antigen levels (HBsAg levels) in IU/ml measured in serum. (B) Hepatitis B e antigen levels (HBeAg) in IU/ml measured in serum. (C) Hepatitis B surface antibody levels mIU/ml measured in serum (D) alanine aminotransferase (ALT) activity in mU/ml measured in serum. (E) Hepatitis B core (HB core) antigen expression in liver. (F) Hepatitis B sAg expression in liver. Dotted black line represents the timing of treatment. Dotted blue line represent the lower limit of quantification (LLOQ) of the assay.

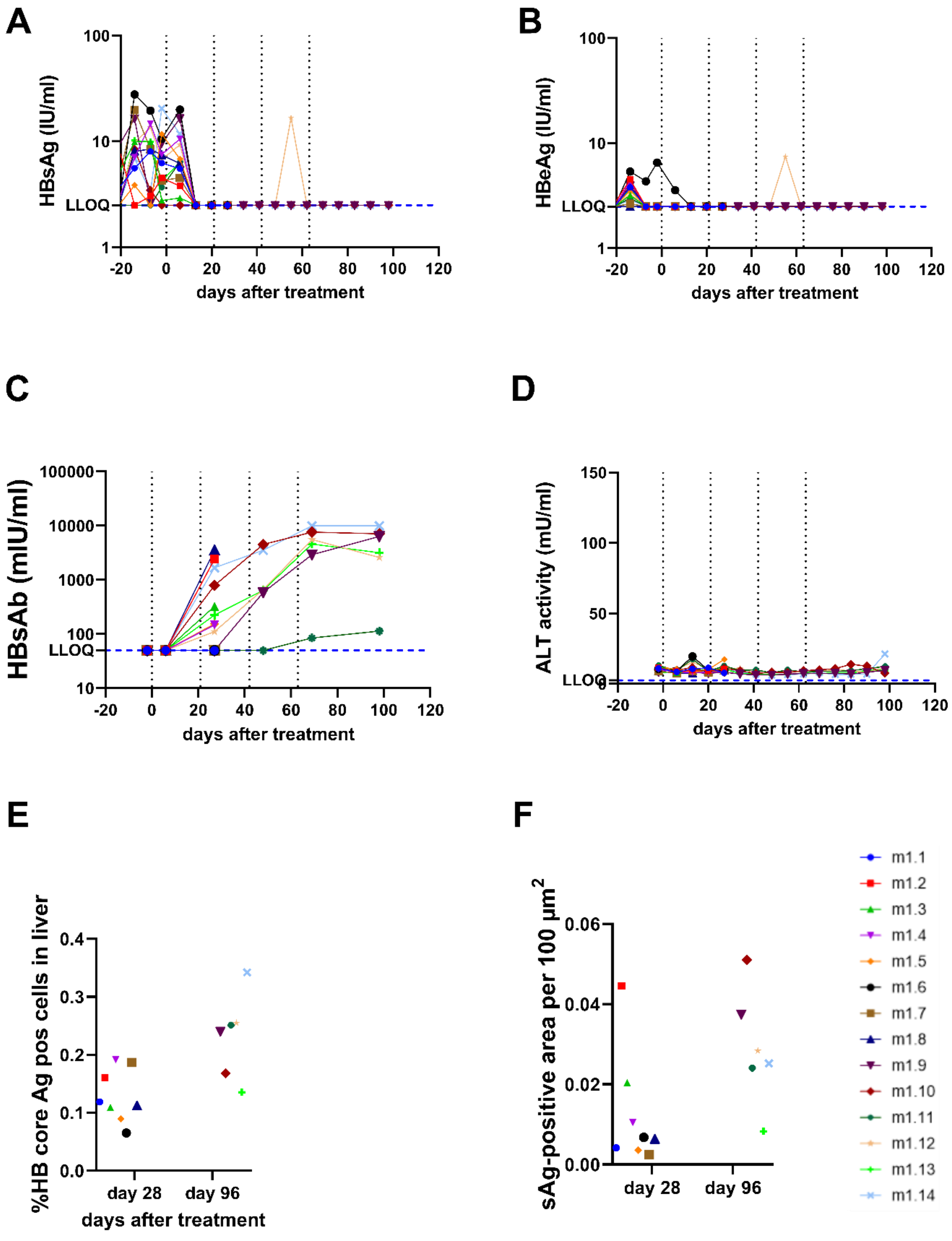

**Figure S2- Viral parameters over time of individual mice that were initially transduced with AAV-HBV ($1x10^9$vge/mice).** Individual-level data are shown; the data correspond to the mean data plotted in Figure 1. (A) Hepatitis B surface antigen levels (HBsAg levels) in IU/ml measured in serum. (B) Hepatitis B e antigen levels (HBeAg) in IU/ml measured in serum. (C) Hepatitis B surface antibody levels mIU/ml measured in serum (D) alanine aminotransferase (ALT) activity in mU/ml measured in serum. (E) Hepatitis B core (HB core) antigen expression in liver. (F) Hepatitis B sAg expression in liver. Dotted black line represents the timing of treatment. Dotted blue line represent the lower limit of quantification (LLOQ) of the assay.

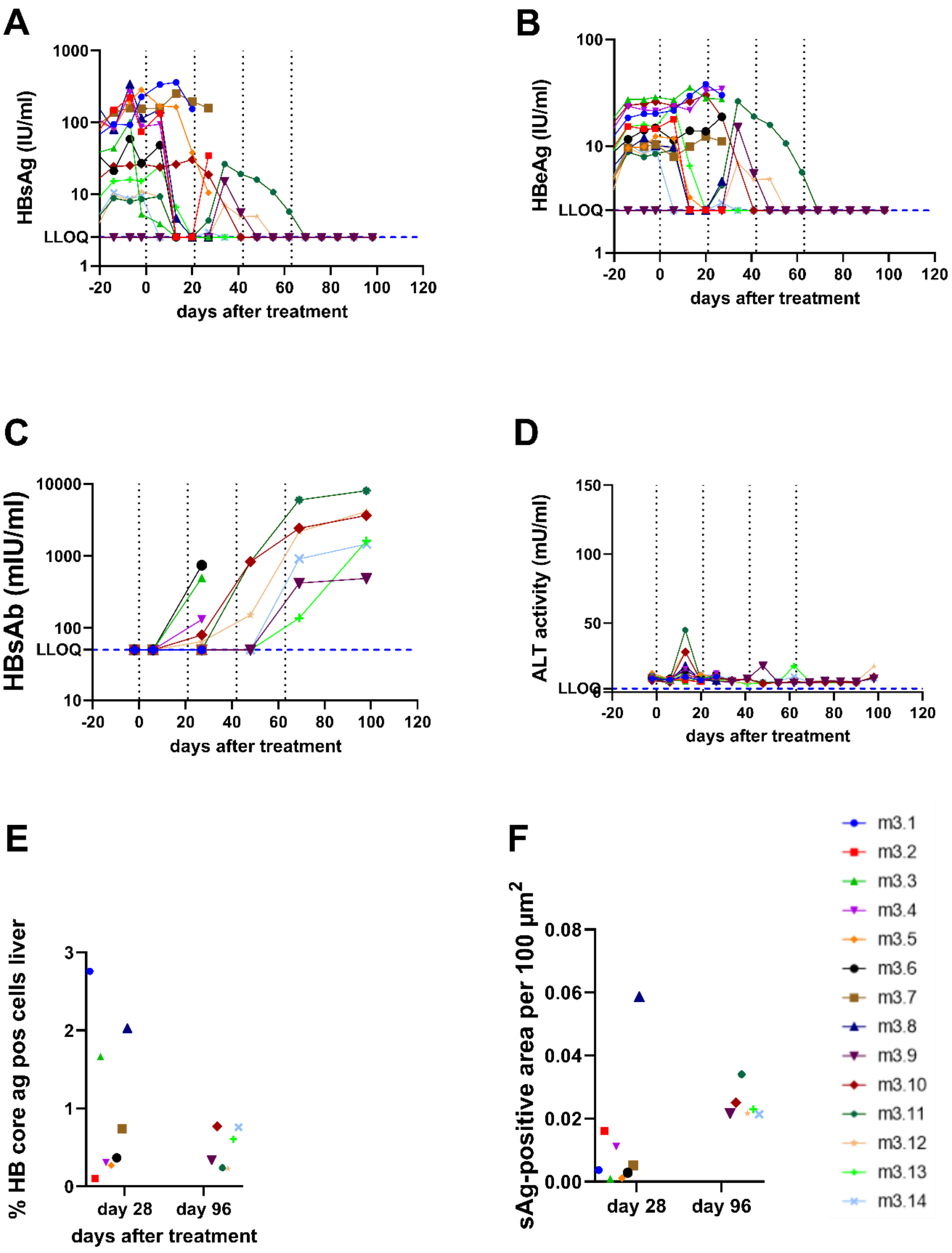

**Figure S3- Viral parameters over time of individual mice that were initially transduced with AAV-HBV (2.5x10$^9$vge/mice).** Individual-level data are shown; the data correspond to the mean data plotted in Figure 1. (A) Hepatitis B surface antigen levels (HBsAg levels) in IU/ml measured in serum. (B) Hepatitis B e antigen levels (HBeAg) in IU/ml measured in serum. (C) Hepatitis B surface antibody levels mIU/ml measured in serum (D) alanine aminotransferase (ALT) activity in mU/ml measured in serum. (E) Hepatitis B core (HB core) antigen expression in liver. (F) Hepatitis B sAg expression in liver. Dotted grey line represents the timing of treatment. Dotted blue line represent the lower limit of quantification (LLOQ) of the assay.

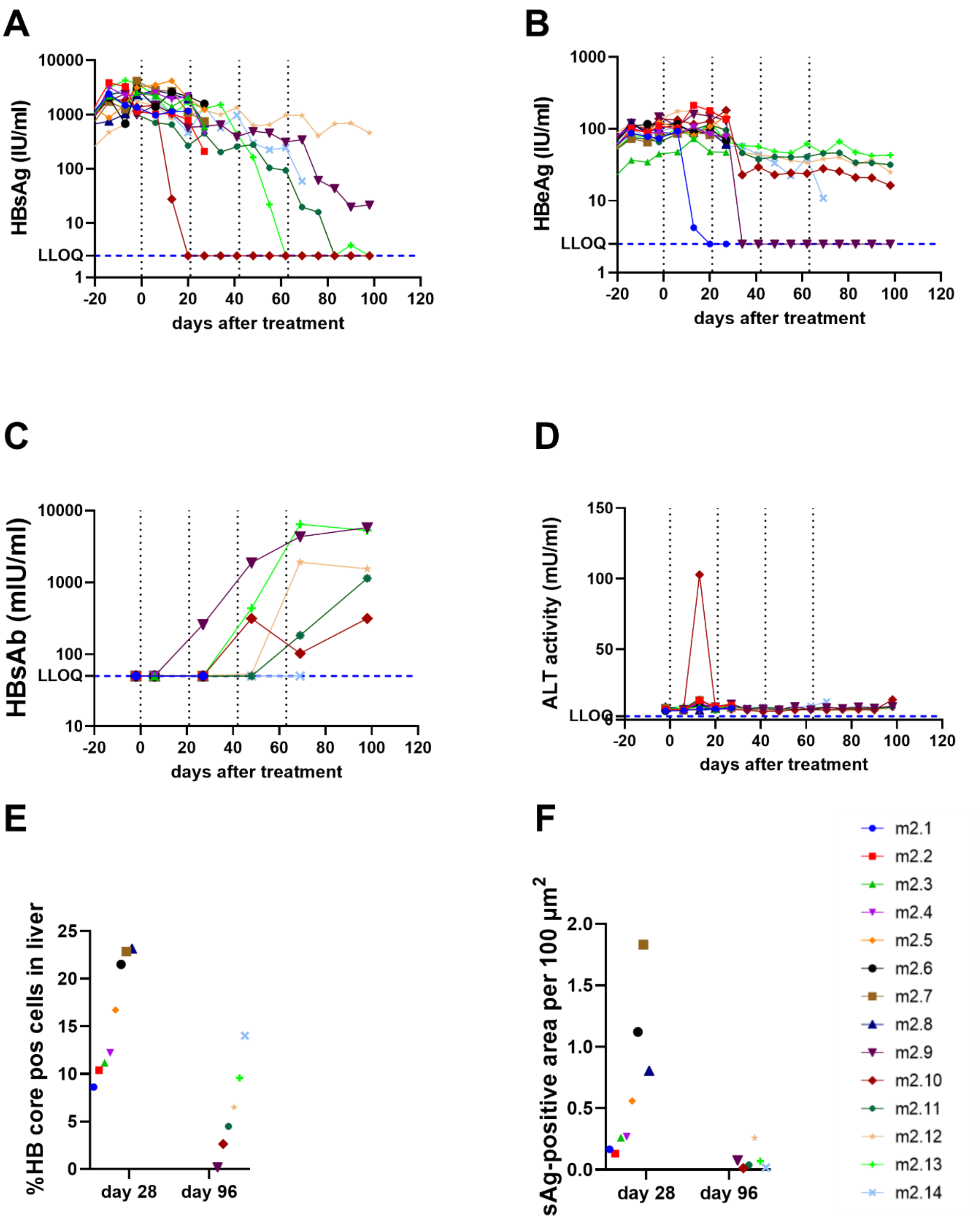

**Figure S4- Viral parameters over time of individual mice that were initially transduced with 2.5x10$^9$vg/ml rAAV-HBV and treated with GalNAc-control siRNA and mock vaccine.** Individual-level data are shown; the data corresponds to the mean data plotted in Figure 3. (A) Hepatitis B surface antigen levels (HBsAg levels) in IU/ml measured in serum until day 70. (B) HBsAg levels in IU/ml measured in serum until day 154. (C) Hepatitis B e antigen levels (HBeAg) in IU/ml measured in serum until day 70. (D) HBeAg levels in IU/ml measured in serum until day 154. (E) Hepatitis B surface antibody levels mIU/ml measured in serum until day 70 (F) HBsAb levels in mIU/ml measured in serum until day 154. (G) alanine aminotransferase (ALT) activity in mU/ml measured in serum until day 70. (H) ALT levels measured in serum until day 154.

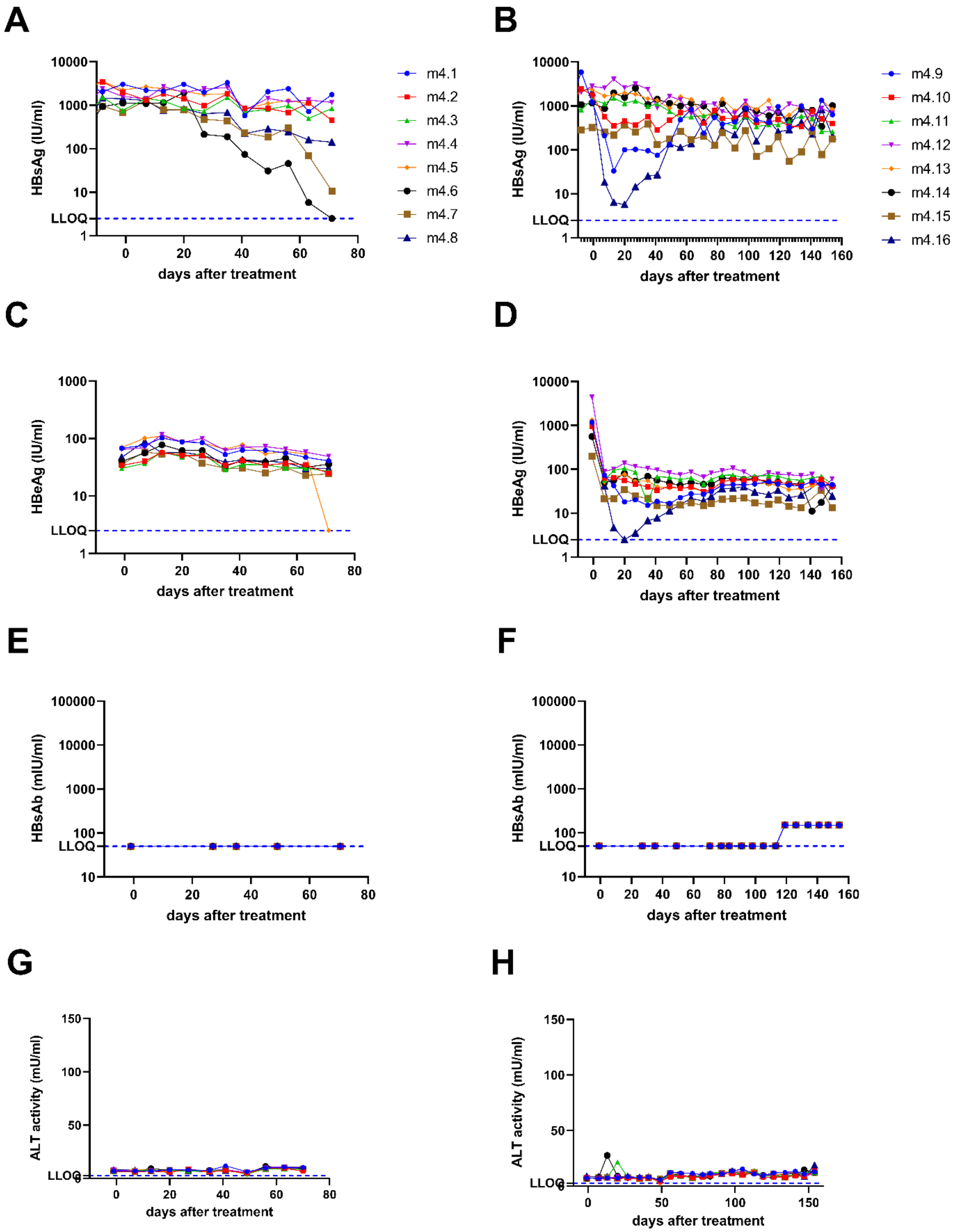

**Figure S5- Viral parameters over time of individual mice that were initially transduced with AAV-HBV (2.5x10$^9$vg/ml) and treated with GalNAc-HBV siRNA and therapeutic vaccine (TxVx).** Individual-level data are shown; the data corresponds to the mean data plotted in Figure 3. (A) Hepatitis B surface antigen levels (HBsAg levels) in IU/ml measured in serum until day 70. (B) HBsAg levels in IU/ml measured in serum until day 154. (C) Hepatitis B e antigen levels (HBeAg) in IU/ml measured in serum until day 70. (D) HBeAg levels in IU/ml measured in serum until day 154. (E) Hepatitis B surface antibody levels mIU/ml measured in serum until day 70 (F) HBsAb levels in mIU/ml measured in serum until day 154. (G) alanine aminotransferase (ALT) activity in mU/ml measured in serum until day 70. (H) ALT levels measured in serum until day 154. Dotted blue line represent the lower limit of quantification (LLOQ) of the assay.

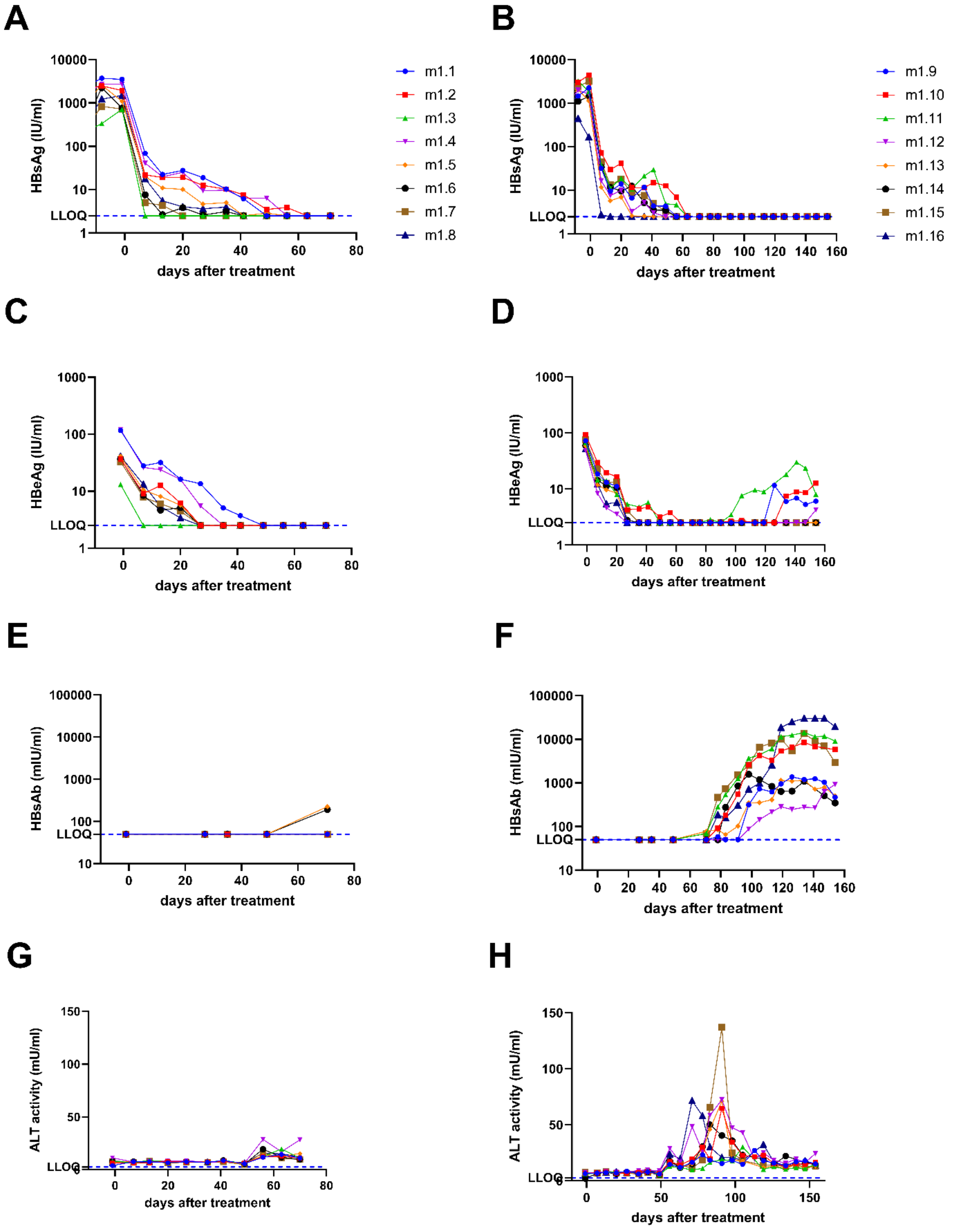

**Figure S6- Viral parameters over time of individual mice that were initially transduced with 2.5x10$^{9}$vg/ml rAAV-HBV and treated with GalNAc-HBV siRNA and mock vaccine.** Individual-level data are shown; the data corresponds to the mean data plotted in Figure 3. (A) Hepatitis B surface antigen levels (HBsAg levels) in IU/ml measured in serum until day 70. (B) HBsAg levels in IU/ml measured in serum until day 154. (C) Hepatitis B e antigen levels (HBeAg) in IU/ml measured in serum until day 70. (D) HBeAg levels in IU/ml measured in serum until day 154. (E) Hepatitis B surface antibody levels mIU/ml measured in serum until day 70 (F) HBsAb levels in mIU/ml measured in serum until day 154. (G) alanine aminotransferase (ALT) activity in mU/ml measured in serum until day 70. (H) ALT levels measured in serum until day 154.

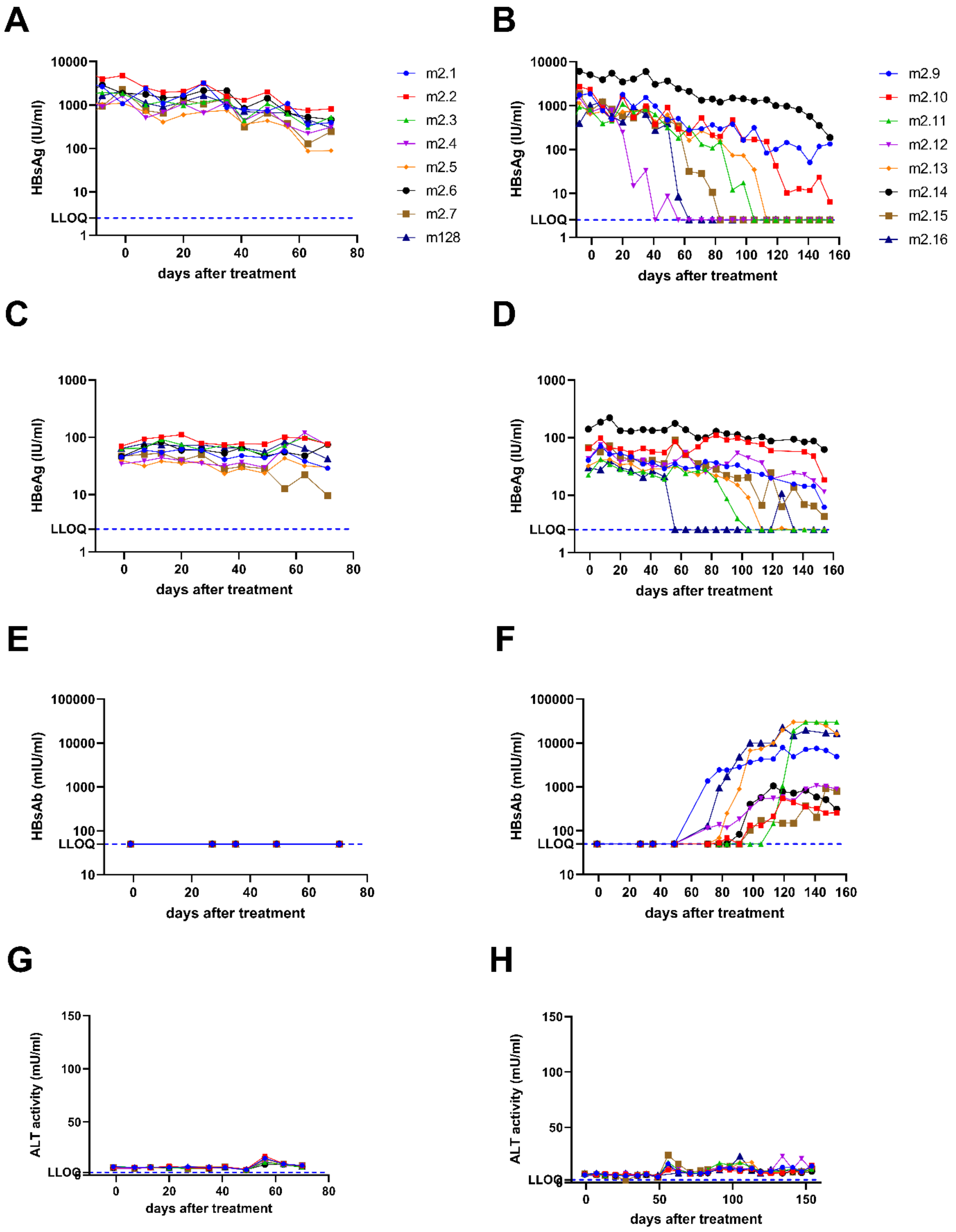

**Figure S7- Viral parameters over time of individual mice that were initially transduced with $2.5x10^9$vg/ml rAAV-HBV and treated with GalNAc-control siRNA and therapeutic vaccine.** Individual-level data are shown; the data corresponds to the mean data plotted in Figure 3. (A) Hepatitis B surface antigen levels (HBsAg levels) in IU/ml measured in serum until day 70. (B) HBsAg levels in IU/ml measured in serum until day 154. (C) Hepatitis B e antigen levels (HBeAg) in IU/ml measured in serum until day 70. (D) HBeAg levels in IU/ml measured in serum until day 154. (E) Hepatitis B surface antibody levels mIU/ml measured in serum until day 70 (F) HBsAb levels in mIU/ml measured in serum until day 154. (G) alanine aminotransferase (ALT) activity in mU/ml measured in serum until day 70. (H) ALT levels measured in serum until day 154

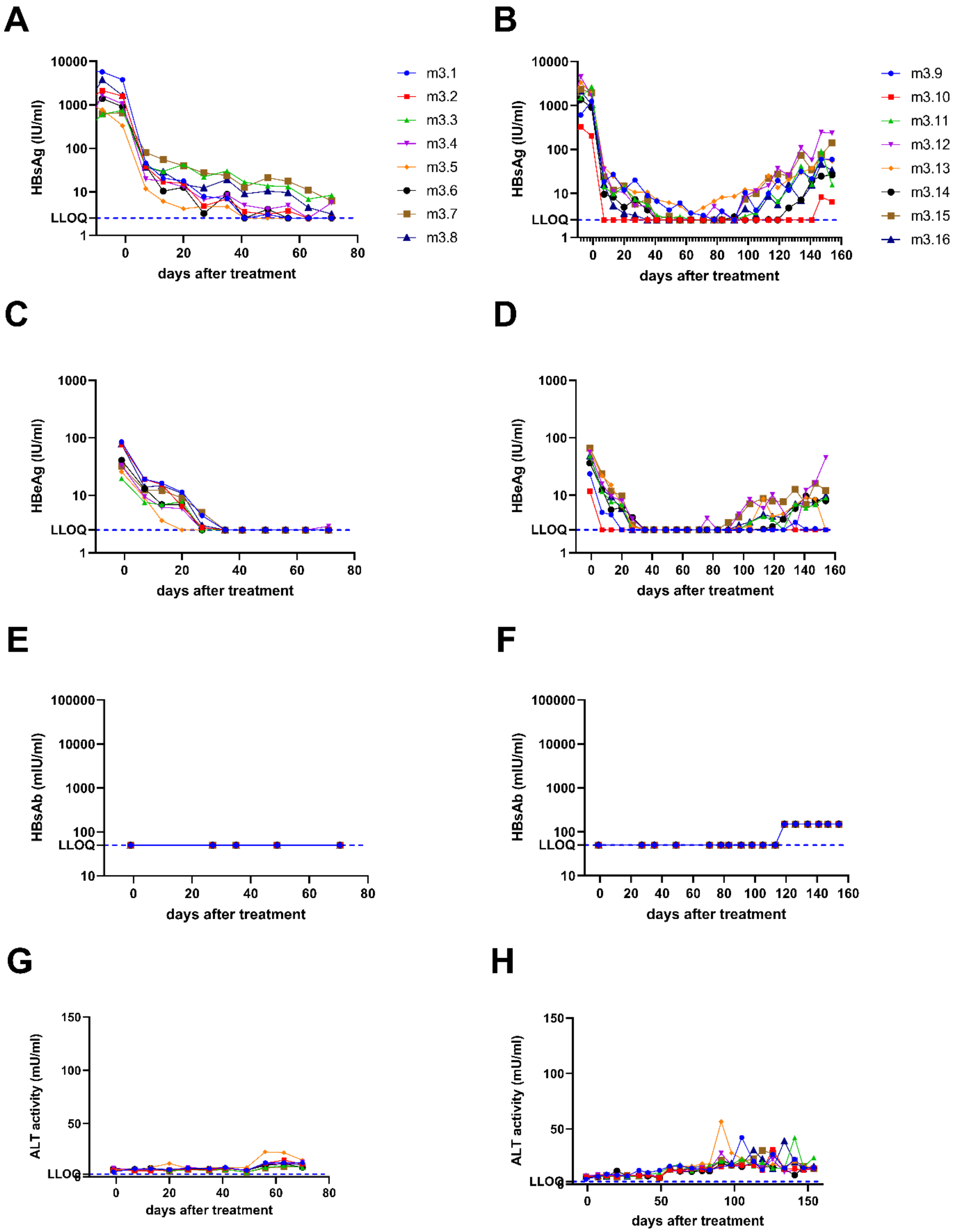

**Figure S8- Single-cell RNA sequencing QC and annotation.** AAV-HBV transduced C57BL/6 mice were treated with GalNAc-HBV siRNA and therapeutic vaccine (G1), with GalNAc-control siRNA and therapeutic vaccine (G2) and with GalNAc-control siRNA and mock vaccine (G3). Intrahepatic immune cells from liver were isolated 1 week after the administration of the second therapeutic vaccine (timepoint day 70) or 7 weeks after the 4th therapeutic vaccine (timepoint day 154). **(a)** Number of cells after QC per sample **(b)** Dendrogram of the levels of cell annotation and cell numbers.

**a**

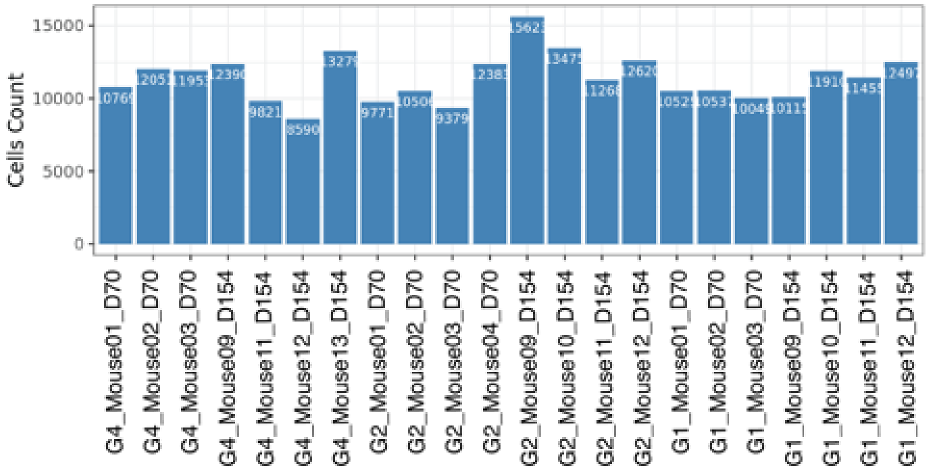


**b**

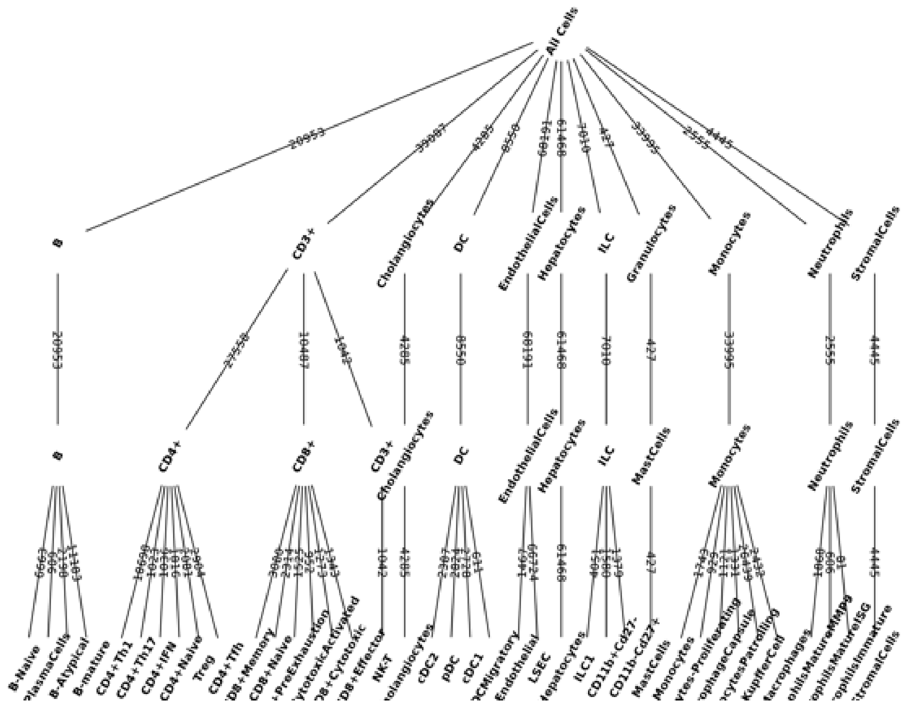

**Figure S9 – CD8 T cell frequency and pre-exhaustion profiles.** AAV-HBV transduced C57BL/6 mice were treated with GalNAc-HBV siRNA and therapeutic vaccine (G1), with GalNAc-control siRNA and therapeutic vaccine (G2) and with GalNAc-control siRNA and mock vaccine (G3). Intrahepatic immune cells from liver were isolated 1 week after the administration of the second therapeutic vaccine (timepoint day 70) or 7 weeks after the 4$^{th}$ therapeutic vaccine (timepoint day 154). **(a)** Frequency of CD8 T cells across groups and statistical comparison at day 154 (paired wilcox test, non-parametric) with fdr adjustment. CD8+Cytotoxic at day 154 from G1 vs G4 was significant (p=0.029, p.adj=0.087). CD8+PreExhaustion at day 154 from G1 vs G4 was significant (p=0.029, p.adj=0.0855). CD8+CentralMemory at day 154 from G1 vs G4 was significant (p=0.029, p.adj=0.0435) and from G1 vs G2 (p=0.029, p.adj=0.0435). CD8+CytotoxicActivated at day 154 from G1 vs G4 was significant (p=0.026, p.adj=0.039) and from G1 vs G2 (p=0.026, p.adj=0.039). **(b)** CD8 pre-exhaustion T cells dot plot of scaled expression of Tcf7 and Id3 across the different groups.

**a**

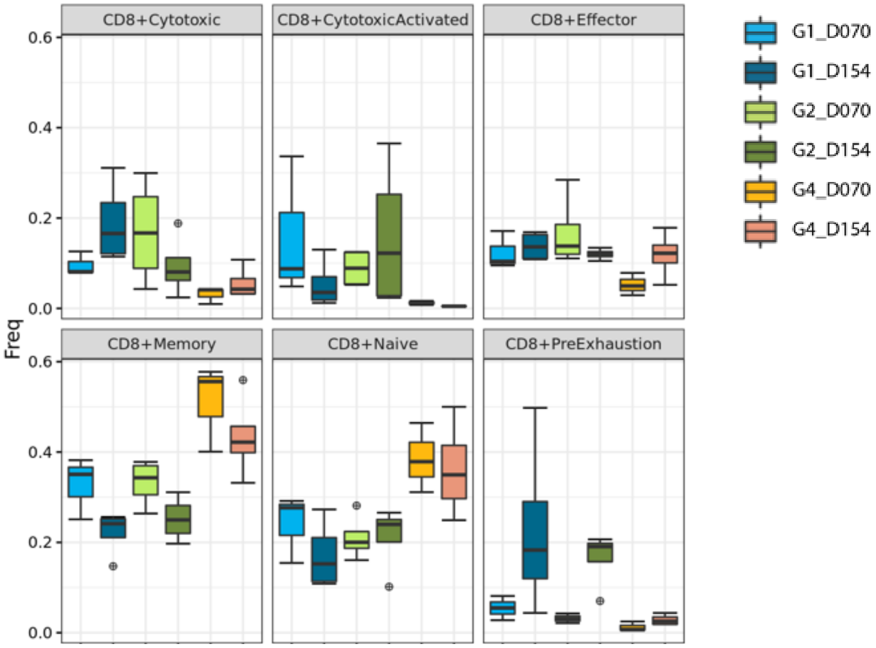
CD8+Cytotoxic
CD8+CytotoxicActivated
CD8+Effector
CD8+Memory
CD8+Naive
CD8+PreExhaustion
Freq
0.0
0.2
0.4
0.6
G1_D070
G1_D154
G2_D070
G2_D154
G4_D070
G4_D154

**b**

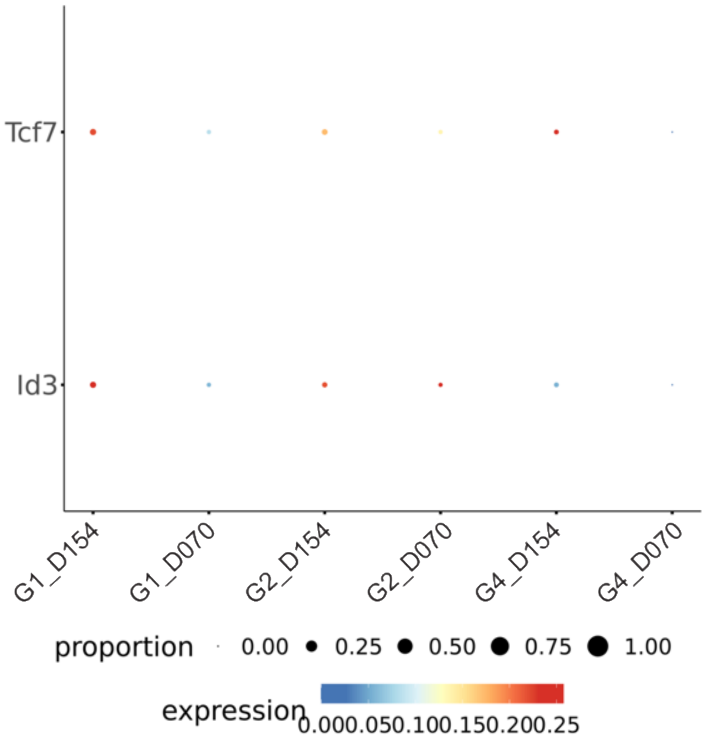
Tcf7
Id3
G1_D154
G1_D070
G2_D154
G2_D070
G4_D154
G4_D070
proportion
0.00
0.25
0.50
0.75
1.00
expression
0.000.050.100.150.200.25

**Fig. S10 – Monocyte and DC compartment frequency analysis.** AAV-HBV transduced C57BL/6 mice were treated with GalNAc-HBV siRNA and therapeutic vaccine (G1), with GalNAc-control siRNA and therapeutic vaccine (G2) and with GalNAc-control siRNA and mock vaccine (G3). Intrahepatic immune cells from liver were isolated 1 week after the administration of the second therapeutic vaccine (timepoint day 70) or 7 weeks after the 4$^{th}$ therapeutic vaccine (timepoint day 154). **(a)** Dotplot from scaled expressed hallmark genes for the antigen presenting cell and monocyte compartments **(b)** Frequency of Monocytes/Macrophages across groups and statistical comparison at day 154 (paired wilcox test, non-parametric, fdr adjustment). Macrophage Capsule at day 154 from G1 vs G4 was significant (p=0.029, p.adj=0.087). Macrophages at day 154 from G1 vs G4 were significant (p=0.029, p.adj=0.0435) and from G1 vs G2 (p=0.029, p.adj=0.0435). **(c)** Frequency of dendritic cells across groups and statistical comparison at day 154 (paired wilcox test, non-parametric, fdr adjustment). cDC1 and pDCs at day 154 from G1 vs G2 were significant (p=0.029, p.adj=0.087). Migratory DCs at day 154 from G1 vs G4 were significant (p=0.029, p.adj=0.057).

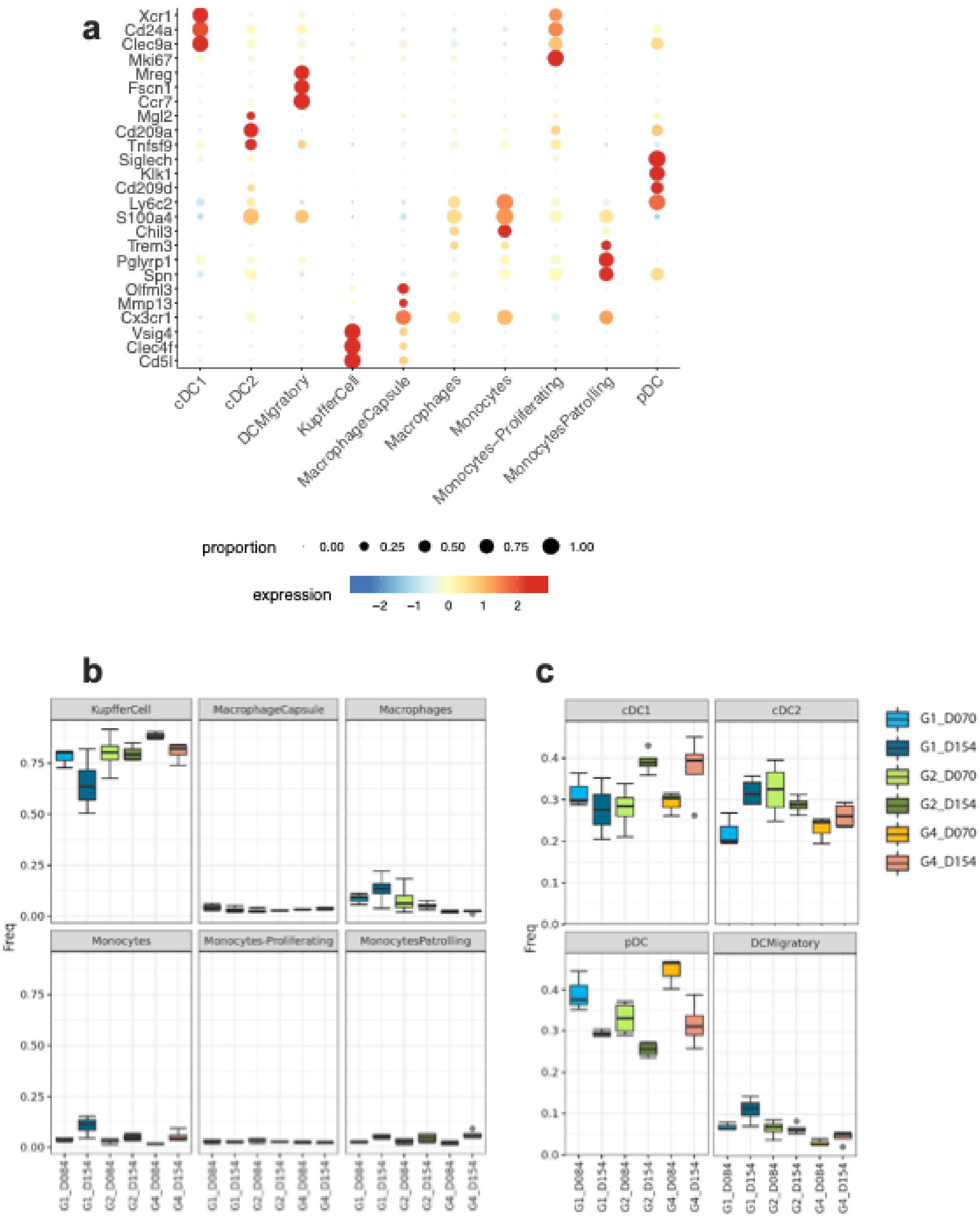

a
Xcr1
Cd24a
Clec9a
Mki67
Mreg
Fscn1
Ccr7
Mgl2
Cd209a
Tnfsf9
Siglech
Klk1
Cd209d
Ly6c2
S100a4
Chil3
Trem3
Pglyrp1
Spn
Olfml3
Mmp13
Cx3cr1
Vsig4
Clec4f
Cd5l
cDC1
cDC2
DCMigratory
KupfferCell
MacrophageCapsule
Macrophages
Monocytes
Monocytes-Proliferating
MonocytesPatrolling
pDC
proportion
0.00
0.25
0.50
0.75
1.00
expression
−2
−1
0
1
2
b
KupfferCell
MacrophageCapsule
Macrophages
Monocytes
Monocytes-Proliferating
MonocytesPatrolling
Freq
0.75
0.50
0.25
0.00
G1_D084
G1_D154
G2_D084
G2_D154
G4_D084
G4_D154
c
cDC1
cDC2
pDC
DCMigratory
Freq
0.4
0.3
0.2
0.1
0.0
G1_D070
G1_D154
G2_D070
G2_D154
G4_D070
G4_D154

**Fig. S11 – ILC compartment frequency analysis.** AAV-HBV transduced C57BL/6 mice were treated with GalNAc-HBV siRNA and therapeutic vaccine (G1), with GalNAc-control siRNA and therapeutic vaccine (G2) and with GalNAc-control siRNA and mock vaccine (G3). Intrahepatic immune cells from liver were isolated 1 week after the administration of the second therapeutic vaccine (timepoint day 70) or 7 weeks after the 4$^{th}$ therapeutic vaccine (timepoint day 154). **(a)** Frequency of ILCs across groups and statistical comparison at day 154 (paired wilcox test, non-parametric) with fdr adjustment. NK_CD11b+Cd27-at day 154 from G1 vs G4 were significant (p=0.029, p.adj=0.087). **(b)** Dot plot with scaled expression of hallmark genes in the ILC compartment.

a

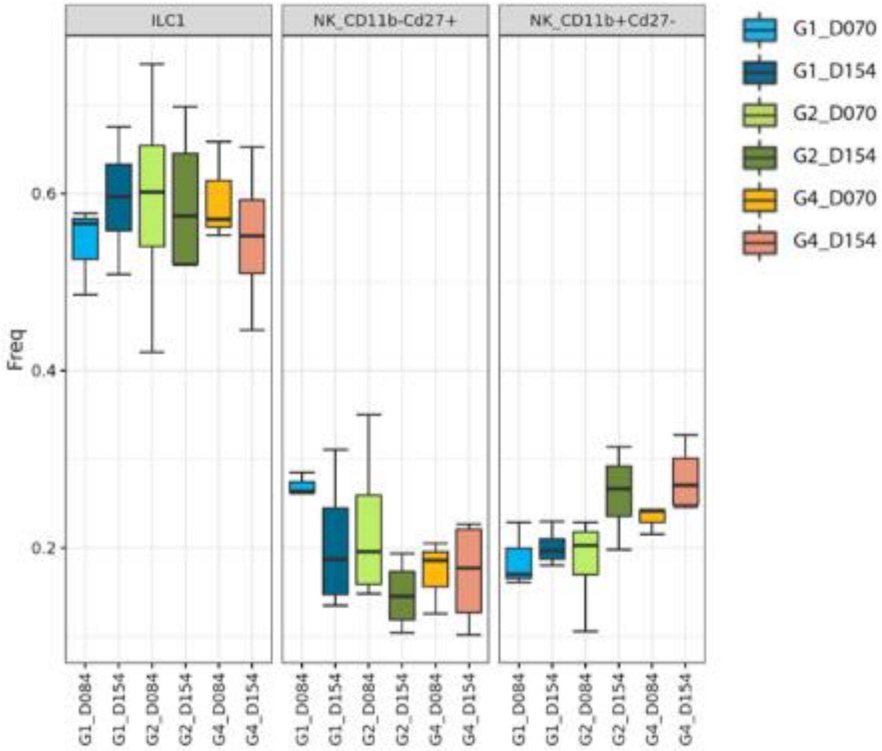
ILC1
NK_CD11b-Cd27+
NK_CD11b+Cd27-
Freq
0.6
0.4
0.2
G1_D084
G1_D154
G2_D084
G2_D154
G4_D084
G4_D154
G1_D070
G1_D154
G2_D070
G2_D154
G4_D070
G4_D154

b

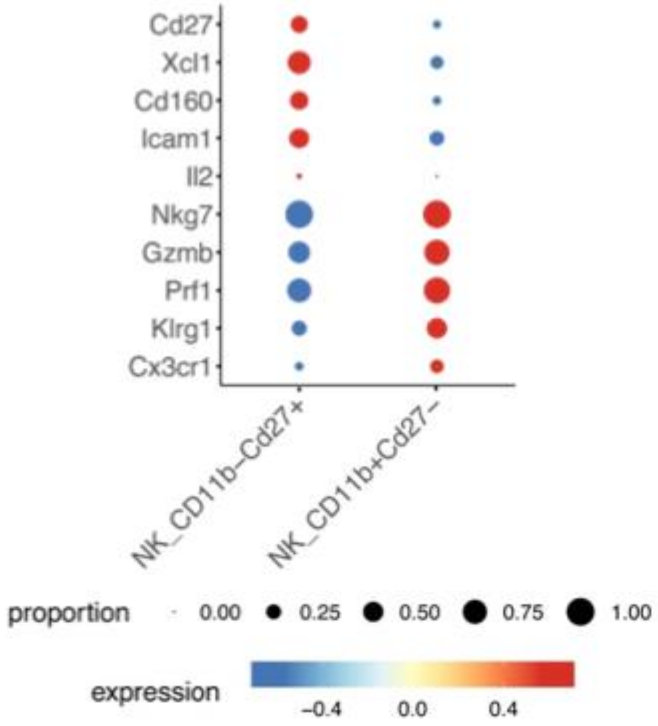
Cd27
Xcl1
Cd160
Icam1
Il2
Nkg7
Gzmb
Prf1
Klrg1
Cx3cr1
NK_CD11b–Cd27+
NK_CD11b+Cd27–
proportion
0.00
0.25
0.50
0.75
1.00
expression
−0.4
0.0
0.4

**Figure S12 – Neutrophil compartment analysis.** AAV-HBV transduced C57BL/6 mice were treated with GalNAc-HBV siRNA and therapeutic vaccine (G1), with GalNAc-control siRNA and therapeutic vaccine (G2) and with GalNAc-control siRNA and mock vaccine (G3). Intrahepatic immune cells from liver were isolated 1 week after the administration of the second therapeutic vaccine (timepoint day 70) or 7 weeks after the 4$^{th}$ therapeutic vaccine (timepoint day 154). **(a)** Dotplot of scaled expression of hallmark genes in the neutrophil compartment. **(b)** Violin plots of normalized expression of IFN/activation genes (*Isg15, Cd274, Rsad2*) and immunosuppressive genes (*Arg1, Mmp9, Mmp8*) **(c)** Frequency of neutrophils across groups and statistical comparison at day 154 (paired wilcox test, non-parametric) with fdr adjustment showed no significant comparisons.

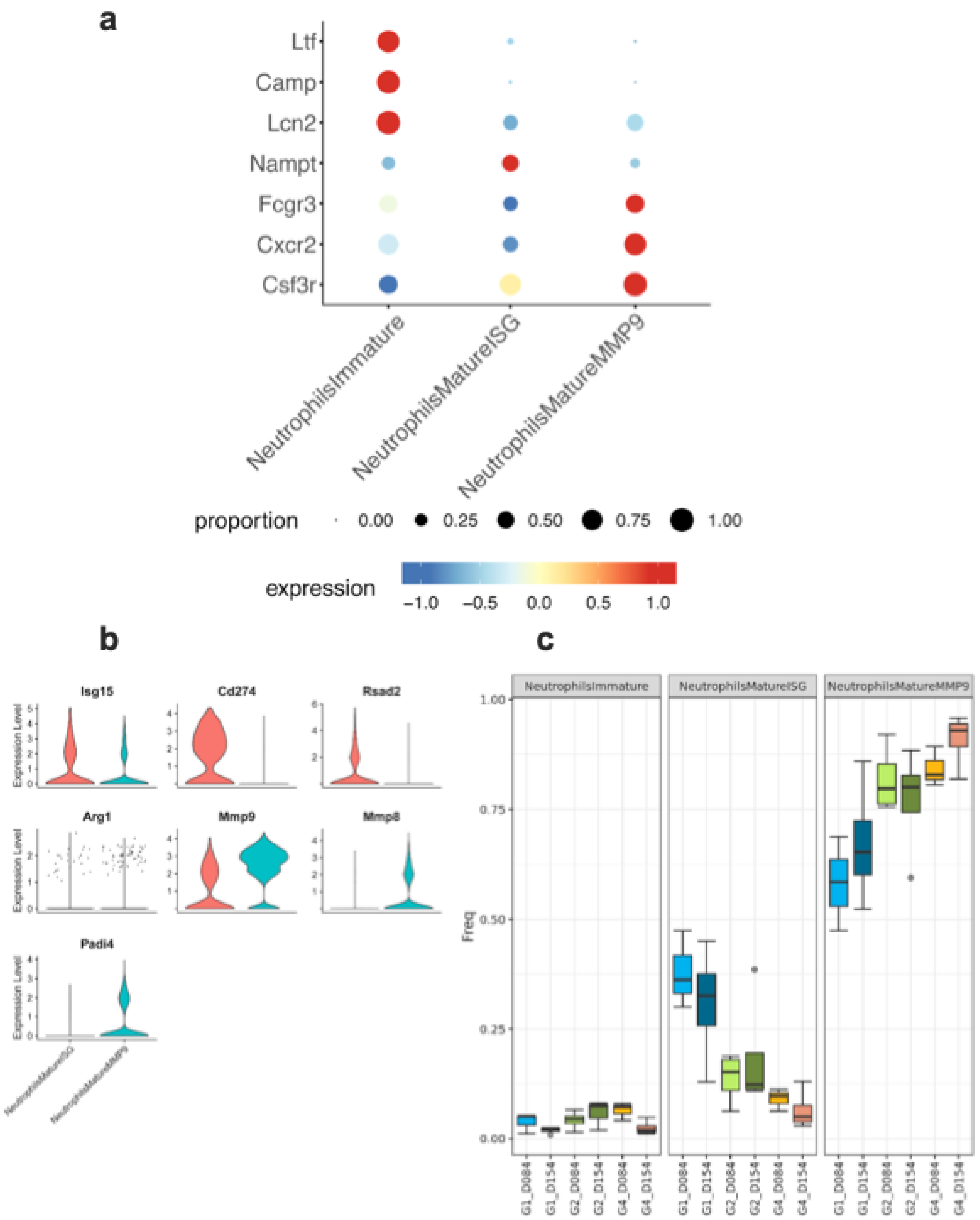
a
Ltf
Camp
Lcn2
Nampt
Fcgr3
Cxcr2
Csf3r
NeutrophilsImmature
NeutrophilsMatureISG
NeutrophilsMatureMMP9
proportion
0.00
0.25
0.50
0.75
1.00
expression
−1.0
−0.5
0.0
0.5
1.0
b
Isg15
Cd274
Rsad2
Arg1
Mmp9
Mmp8
Padi4
Expression Level
c
NeutrophilsImmature
NeutrophilsMatureISG
NeutrophilsMatureMMP9
Freq
1.00
0.75
0.50
0.25
0.00
G1_D084
G1_D154
G2_D084
G2_D154
G4_D084
G4_D154

**Figure S13 –TCR clonality frequency** AAV-HBV transduced C57BL/6 mice were treated with GalNAc-HBV siRNA and therapeutic vaccine (G1), with GalNAc-control siRNA and therapeutic vaccine (G2) and with GalNAc-control siRNA and mock vaccine (G3). Intrahepatic immune cells from liver were isolated 1 week after the administration of the second therapeutic vaccine (timepoint day 70) or 7 weeks after the 4th therapeutic vaccine (timepoint day 154). **(a)** Frequency of the TCR clonal expansion in the CD4 compartment per group **(b)** Frequency of the TCR clonal expansion in the CD8 compartment per group.

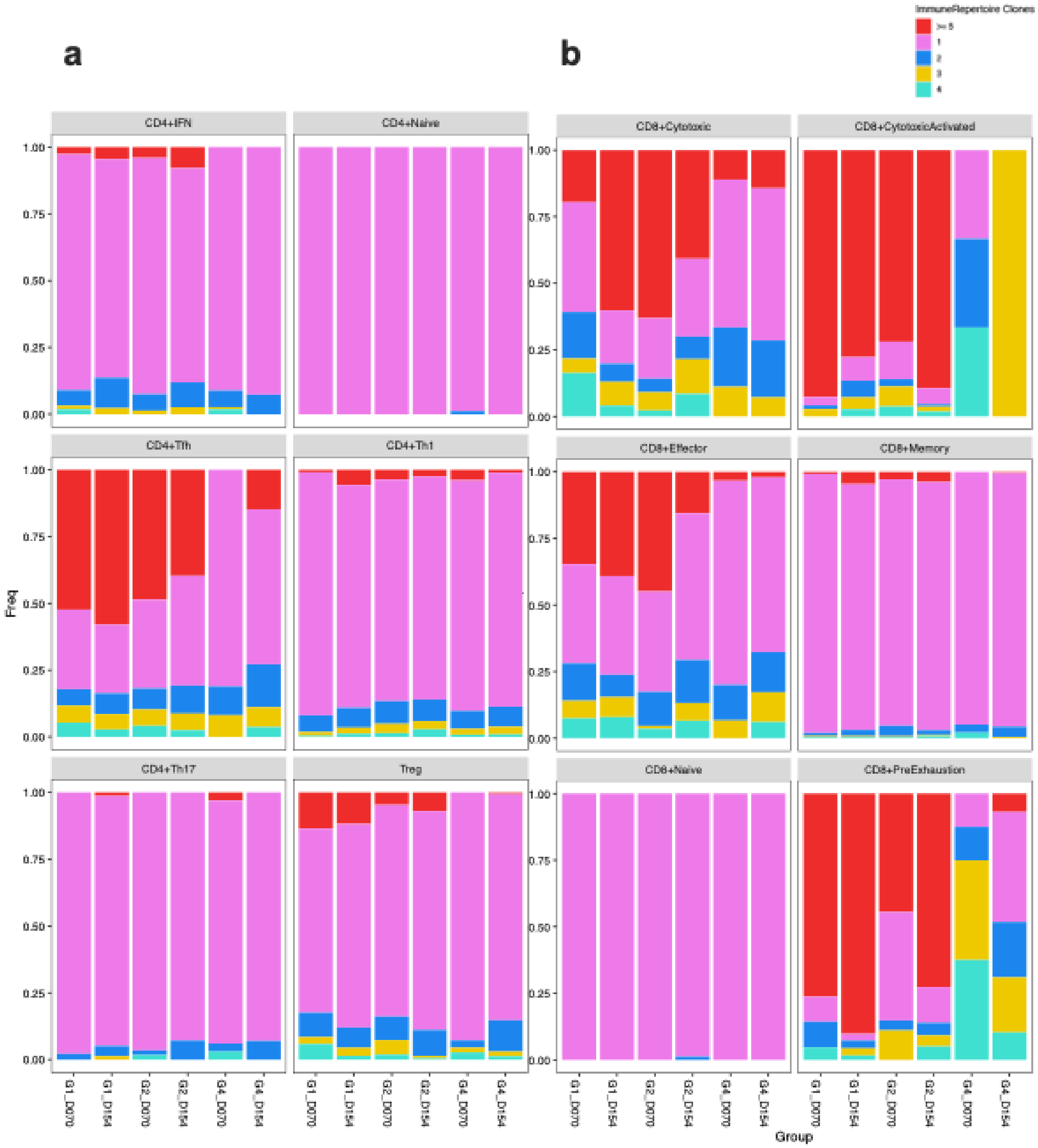

**Figure S14 – B cell compartment analysis.** AAV-HBV transduced C57BL/6 mice were treated with GalNAc-HBV siRNA and therapeutic vaccine (G1), with GalNAc-control siRNA and therapeutic vaccine (G2) and with GalNAc-control siRNA and mock vaccine (G3). Intrahepatic immune cells from liver were isolated 1 week after the administration of the second therapeutic vaccine (timepoint day 70) or 7 weeks after the 4$^{th}$ therapeutic vaccine (timepoint day 154). **(a)** Violin plots of normalized expression from B-cell marker genes for each cell subtype **(b)** Frequency of the BCR clonal expansion in each B cell subtype, in red are depicted the expanded clonotypes. **(c)** Dotplot of scaled expression of Long-Lived Plasma Cell marker genes in Plasma cells, across clonality groups, >=3 clonotypes are expanded. **(d)** Phylogeny analysis from B cell clones (CDR3 sequences) of samples from the siRNA and therapeutic vaccine combination **(e)** from the control siRNA and therapeutic vaccine treatment **(f)** and from the control siRNA and empty plasmid.

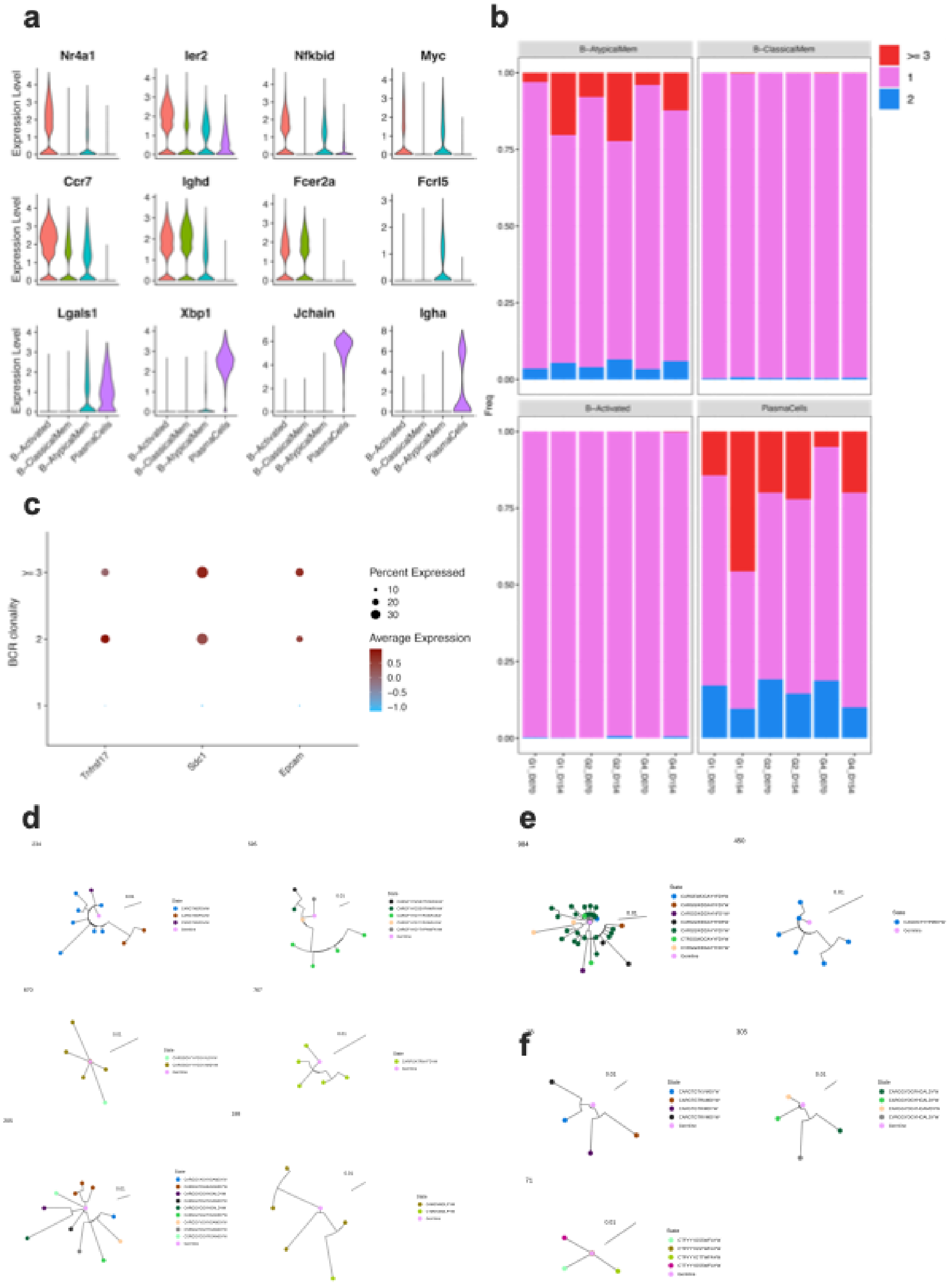
a
Nr4a1
Ier2
Nfkbid
Myc
Ccr7
Ighd
Fcer2a
Fcrl5
Lgals1
Xbp1
Jchain
Igha
Expression Level
B-Activated
B-ClassicalMem
B-AtypicalMem
PlasmaCells
b
B-AtypicalMem
B-ClassicalMem
B-Activated
PlasmaCells
Freq
>= 3
1
2
c
BCR clonality
Percent Expressed
Average Expression
Tnfrsf17
Sdc1
Epcam
d
e
f

**Supplementary Table 1 – Percentage distribution of cell types**

| CellType | TotalCells | Percentage |
|---|---|---|
| B | 20953 | 8 |
| CD3+ | 39087 | 16 |
| Cholangiocytes | 4285 | 2 |
| DC | 8550 | 3 |
| EndothelialCells | 68191 | 27 |
| Granulocytes | 427 | 0 |
| Hepatocytes | 61468 | 24 |
| ILC | 7010 | 3 |
| Monocytes | 33995 | 14 |
| Neutrophils | 2555 | 1 |
| StromalCells | 4445 | 2 |
| Totals | 250966 | 100 |
